# Robust T1 MRI cortical surface pipeline for neonatal brain and systematic evaluation using multi-site MRI datasets

**DOI:** 10.1101/2021.01.13.426611

**Authors:** Mengting Liu, Claude Lepage, Sharon Y. Kim, Seun Jeon, Sun Hyung Kim, Julia Pia Simon, Nina Tanaka, Shiyu Yuan, Tasfiya Islam, Bailin Peng, Knarik Arutyunyan, Wesley Surento, Justin Kim, Neda Jahanshad, Martin A. Styner, Arthur W. Toga, A. James Barkovich, Duan Xu, Alan C. Evans, Hosung Kim

**Author notes:** **Corresponding author:** Hosung Kim, PhD, Assistant Professor of Neurology, USC Stevens Neuroimaging and Informatics Institute, Keck School of Medicine of USC, University of Southern California, 2025 Zonal Ave., Los Angeles, CA 90033. Competing interests: None. Sponsorships: None.

## Abstract

The human brain grows the most dramatically during the perinatal and early postnatal periods, during which preterm birth or perinatal injury that may alter brain structure and lead to developmental anomalies. Thus, characterizing cortical thickness of developing brains remains an important goal. However, this task is often complicated by inaccurate cortical surface extraction due to small-size brains. Here, we propose a novel complex framework for the reconstruction of neonatal WM and pial surfaces, accounting for large partial volumes due to small-size brains. The proposed approach relies only on T1-weighted images unlike previous T2-weighted image-based approaches while only T1-weighted images are sometimes available under the different clinical/research setting. Deep neural networks are first introduced to the neonatal MRI pipeline to address the mis-segmentation of brain tissues. Furthermore, this pipeline enhances cortical boundary delineation using combined models of the CSF/GM boundary detection with edge gradient information and a new skeletonization of sulcal folding where no CSF voxels are seen due to the limited resolution. We also proposed a systematic evaluation using three independent datasets comprising 736 preterm and 97 term neonates. Qualitative assessment for reconstructed cortical surfaces shows that 86.9% are rated as accurate across the three site datasets. In addition, our landmark-based evaluation shows that the mean displacement of the cortical surfaces from the true boundaries was less than a voxel size (0.532±0.035mm). Evaluating the proposed pipeline (namely NEOCIVET 2.0) shows the robustness and reproducibility across different sites and different age-groups. The mean cortical thickness measured positively correlated with postmenstrual age (PMA) at scan (p<0.0001); Cingulate cortical areas grew the most rapidly whereas the inferior temporal cortex grew the least rapidly. The range of the cortical thickness measured was biologically congruent (1.3mm at 28 weeks of PMA to 1.8mm at term equivalent). Cortical thickness measured on T1 MRI using NEOCIVET 2.0 was compared with that on T2 using the established dHCP pipeline. It was difficult to conclude that either T1 or T2 imaging is more ideal to construct cortical surfaces. NEOCIVET 2.0 has been open to the public through CBRAIN (https://mcin-cnim.ca/technology/cbrain/), a web-based platform for processing brain imaging data.

## Introduction

The human brain grows the most dramatically during the perinatal and postnatal periods (Knickmeyer et al., 2008), where the majority of cortical growth and folding partition the brain for various sophisticated functions (Alexander-Bloch et al., 2018). This critical time window of neurodevelopment is susceptible to developmental anomalies, such as preterm birth or perinatal injuries. The effect of such impairments on early development, although documented through limited histological or qualitative clinical imaging studies (Battin et al., 1998;Kapellou et al., 2006;Ball et al., 2012;Ball et al., 2013;Pandit et al., 2014;Ball et al., 2015;Guo et al., 2016;Lefevre et al., 2016;Kim et al., 2020), still remain far from being completely understood. Given vast gaps in knowledge of early brain development, it remains a paramount goal to accurately measure cortical morphological changes in neonates in order to elucidate post-natal brain development and neurodevelopmental outcomes in preterm survivors (Habas et al., 2012;Wright et al., 2014).

Modern neuroimaging approaches have provided the ability to more accurately and non-invasively characterize and study brain morphology, such as through *in vivo* T1-weighted (T1w) and T2-weighted (T2w) magnetic resonance imaging (MRI). However, unlike MRI methods implemented to study the adult brain, MRI methods and processing frameworks for the neonatal brain are still under investigation in order to circumvent substantial challenges, such as opposite/reduced tissue contrast between WM and GM, large within-tissue intensity variations, and regionally-heterogeneous image appearances that dynamically change in accordance with neural development. In particular, these challenges lead to significant difficulties in quantifying cortical morphology measurements, including cortical thickness, surface curvature, and sulcal depth, which demand accurate surface reconstruction methods. Conventional pipelines such as CIVET (MacDonald et al., 2000;Kim et al., 2005), Freesurfer (Fischl, 2012), and Caret (Van Essen et al., 2001) are not suitable for neonatal brains, as they rely on established tissue contrast normally exhibited in adult brain MRIs. As such, several modifications (Hill et al., 2010;Leroy et al., 2011;Wright et al., 2015;Kim et al., 2016) have been incorporated into such pipelines to optimize the surface reconstruction of neonatal brains. However, these methods provide only the successful reconstruction of the inner cortical surface, while remaining incapable of robustly measuring cortical thickness, which is a crucial morphological characteristic of brain development (Raznahan et al., 2011a).

One of the main challenges in reconstructing the outer cortical border is the significant partial volume effects (Osechinskiy and Kruggel, 2012;Li et al., 2019) located in areas of the brain scan where only scattered cerebrospinal fluid (CSF) voxels are found within the deep and narrow sulcal fundi. To overcome this issue, previous studies in adult brain MRI mainly included one of the two following strategies: (1) generate a skeleton representing the medial surface of each sulcus (Mangin et al., 1995;Han et al., 2001;Han et al., 2004;Kim et al., 2005;Li et al., 2012), or (2) expand the initial cortical surface mesh (placed in the GM/WM border) onto opposing gyral folds towards each other until either GM and CSF reach a boundary or they are close enough to each other (Dale et al., 1999). Such adult-specific methods are based on the assumption that the partial volumes of CSF can be accurately modeled, and the GM/CSF boundary can be clearly identified. However, these assumptions cannot be applied to neonatal brain scans, due to their limited image resolution, and smaller head sizes which exacerbate partial volume effects in sulcal CSF (see Fig. 1).

**Fig 1.**
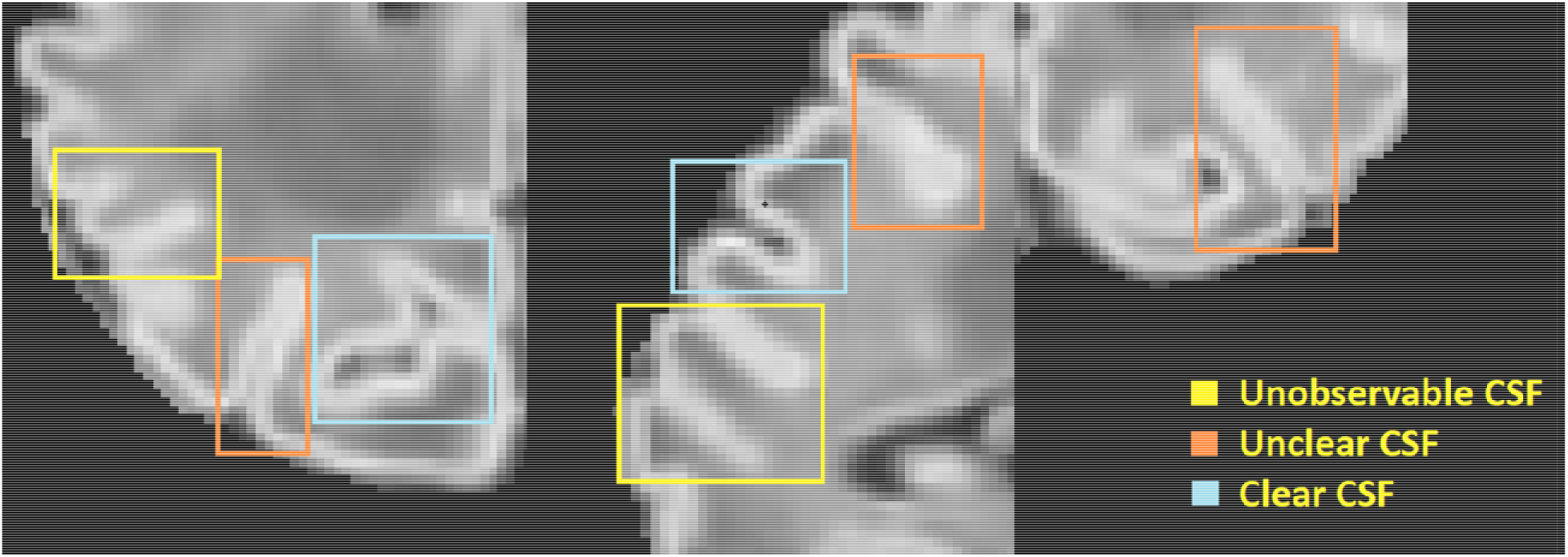
An illustration of large partial volume effect in deep sulcus.

To more accurately generate the outer cortical surface, Li and his colleagues (Li et al., 2014) applied a deformable surface model to longitudinal scans where CSF could be identified on the images acquired from older infants (e.g., 1 year old and older). Such MRIs would be acquired longitudinally in a small number of neonates under a planned research setting. The developing Human Connectome Project (dHCP) pipeline (Schuh et al., 2017;Makropoulos et al., 2018a) proposed the expansion of the inner cortical surface to the outer rim based on a non-intersection constraint and a local edge detection. The partial volume effect was corrected using a draw-EM tissue segmentation approach with specially designed sulci detection and enhancement approach (Makropoulos et al., 2016). However, this surface reconstruction could be performed only when T2w images were available. A more recent pipeline (Zöllei et al., 2020) performed surface reconstruction for 0-2 years old infants using T1w images only. This method used a multiatlas-based segmentation combined with Bayesian inference to solve the fundamental issue of unobservable CSF volumes within a sulcus (Fig. 1). However, when applied to T1w images in the dHCP dataset, the quality of the generated surfaces using this method was much worse compared to those generated using the pipeline conducted on T2w images (Makropoulos et al., 2018b).

Given that volumetric T2-weighted images are not frequently used in infants due to poor contrast (Zöllei et al., 2020), a robust neonatal surface generation pipeline solely dependent on T1w images is needed. In this study, we propose a complete pipeline that robustly reconstructs inner cortical and pial surfaces of neonatal brains, while computing sulcal depth and cortical thicknes measurements. This approach requires only one type of image modality (T1w MRI) and includes an additional number of novel features, compared to our previous pipeline (Kim et al., 2016) and other approaches, that address the aforementioned challenges. First, our new pipeline integrates a deep learning-based tissue segmentation method using nonlocal 3D-Unet into the cortical morphometry framework to address for regionally varying tissue contrast in neonates. Our approach resulted in higher accuracy compared to other neonatal brain tissue segmentation methods (Weisenfeld and Warfield, 2009;Gousias et al., 2013;Wang et al., 2013;Makropoulos et al., 2014;Wang et al., 2015;Beare et al., 2016). Second, CSF volumes are skeletonized by combining CSF-PVs with sulcal GM, which approximately place outer cortical surfaces onto GM/CSF boundaries. Third, our pipeline incorporates the generated CSF skeleton with CSF/GM edges, which are detected by a radial gradient-based approach, in order to more accurately identify candidate locations of the pial surface. Unlike the dHCP pipeline (Schuh et al., 2017;Makropoulos et al., 2018a), where the gradient-based refinement for pial surface was performed by directly deforming the WM surface, the gradient-based refinement for pial surface in our design is performed within a short distance range after the GM/CSF boundary is first approximated and placed at the skeleton. Finally, we systematically perform qualitative and quantitative evaluations using multi-site neonatal datasets to investigate whether the proposed pipeline could overcome the limitations in the modules that might be suboptimal for regional analyses of poor-contrast neonatal cortex. We compute cortical thickness values of over 80,000 vertices sampled on the constructed surface. After registering individual surfaces to their age-matched template, the results suggest that the proposed cortical surface reconstruction can robustly provide measurements that characterize realistic neonatal brain morphological changes across the multi-site datasets.

## Materials and Equipment

### Subjects

Our first dataset comprised of 223 preterm newborns (mean PMA at birth = 28.1 ± 2.0 weeks; range 24–33 weeks), admitted to UCSF Benioff Children’s Hospital San Francisco between June 2006 and January 2016. Exclusion criteria included (i) clinical evidence of a congenital malformation or syndrome, (ii) congenital infection, and (iii) newborns too clinically unstable for transport to and from the MRI scanner. Parental consent was obtained for all cases following a protocol approved by the institutional Committee on Human Research. All patients were scanned postnatally as soon as they became clinically stable (PMA at scan: 31.7 ± 1.8 weeks; range 26– 40 weeks), and 132 patients were re-scanned before discharge at late preterm age (PMA at scan: 35.9±2.0 weeks; range 32–44 weeks). Due to extremely severe motion artifact, 36 baseline and 25 follow-up scans were excluded. The final database included 187 baseline (PMA = 31.8 ± 1.8 weeks) and 107 follow-up scans (36.0 ± 2.0 weeks).

We also utilized two independent datasets collected from the University of Northern Carolina (UNC) and T1w images from the Developing Human Connectome Project (dHCP) to cross-validate our pipeline. The UNC dataset includes 57 term born neonates (PMA = 39.2 ± 1.3 weeks) who were scanned at PMA of 40-45 weeks. The 492 T1w images (PMA = 39.2 ± 3.8 weeks) in the dHCP dataset, which were scanned at PMA of 29-45 weeks, were also included.

### Image acquisition

At UCSF, customized MRI-compatible incubators with specialized head coils were used to provide a quiet, well-monitored environment for neonates during the MRI scan, minimizing patient movement and improving the signal-to-noise ratio. Newborns enrolled between June 2006 and July 2011 (n = 95) were scanned on a 1.5-Tesla General Electric Signa HDxt system (GE Medical systems, Waukesha, WI, USA) using a specialized high-sensitivity neonatal head coil built within a custom-built MRI compatible incubator. T1-weighted images were acquired using sagittal 3-dimensional inversion recovery spoiled gradient echo (3D SPGR) (repetition time [TR] = 35 ms; echo time [TE] = 6.00 ms; inversion time of 0.00 ms; field of view [FOV] = 256 × 192 mm^2^; number of excitations [NEX] = 1.00; and flip angle [FA] = 35°), yielding images with 1 × 1 × 1 mm^3^ spatial resolution. Newborns enrolled between July 2011 and March 2015 (n=128) were scanned on a 3-Tesla General Electric Discovery MR750 system in a different MR compatible incubator and using a specially designed (for 3 T imaging) neonatal head coil. T1-weighted images were also acquired using sagittal 3D IR-SPGR (TR = minimum; TE = minimum; inversion time of 450 ms; FOV = 180 × 180 mm^2^; NEX = 1.00; FA = 15°), and were reformatted in the axial and coronal planes, yielding images with 0.7 × 0.7 × 1 mm^3^ spatial resolution.

For the UNC dataset, the imaging protocol parameters match those used by (Kim et al., 2016b). Neonates were scanned on a 3T Siemens Trio system. T1-weighted images were acquired using 3-dimensional magnetization-prepared radio-frequency pulses and rapid gradient-echo (MP-RAGE; repetition time [TR] = 2400ms; echo time [TE] = 3.16ms; inversion time [TI] = 1200ms; and flip angle [FA] = 8°), yielding images with 1 × 1 × 1 mm^3^ spatial resolution.

## Methods

### Pipeline overview

Our new framework has extended the original NEOCIVET pipeline (henceforth NEOCIVET 1.0) (Kim et al., 2016) to include the reconstruction of neonatal pial surfaces (GM-CSF interface) as well as the improvement of existing features. The new pipeline begins with general data pre-processing, including denoising and intensity nonuniformity correction. Then, the brain is extracted using a deep 3D-Unet (Hwang et al., 2019) and registered to the MNI-NIH neonatal brain template (http://www.bic.mni.mcgill.ca/ServicesAtlases/NIHPD-obj2), with spatial resolution of 0.6 × 0.6 × 0.6 mm^3^ (isotropic). Different types of brain tissue (GM, WM, and CSF) are thereafter segmented by an advanced deep non-local 3D-Unet (Wang et al., 2020b). Individual templates (MRI + manually segmented tissue labels) utilized for the deep learning approach are then selected evenly across all postmenstrual ages (PMA). Next, the corpus callosum is segmented on the midline-plane and used to divide the WM into hemispheres. Instead of the original parametric deformable model, a marching-cube based framework is newly adopted to generate a triangulated mesh WM surface attached to the boundary between the GM and WM. This process results in the number of surface-meshes adapted to the individually varying brain size and folding area, addressing the issue of the complex optimization of the parameters needed in the previous parametric model (Kim et al., 2016). After resampling to a fixed number of 81,920 surface meshes (triangles) using the icosahedron spherical fitting, this surface is further fitted to the sharp edge of the GM-WM interface based on the image intensity gradient, which preserves the spherical topology of the cortical mantle. A CSF skeleton is then generated from the union of GM and CSFs. Pial surface is constructed by expanding the WM surface towards the skeleton as an intermediate pial surface. The intermediate pial surface further undergoes a fine deformation to identify actual edges of sulcal CSF volumes using an intensity gradient feature model. Finally, the cortical thickness is estimated based on the shortest distance between the white matter and pial surface, with a smooth kernel size of 10 mm (Vasung et al., 2016). Fig. 2 presents the workflow of our proposed pipeline.

**Fig 2.**
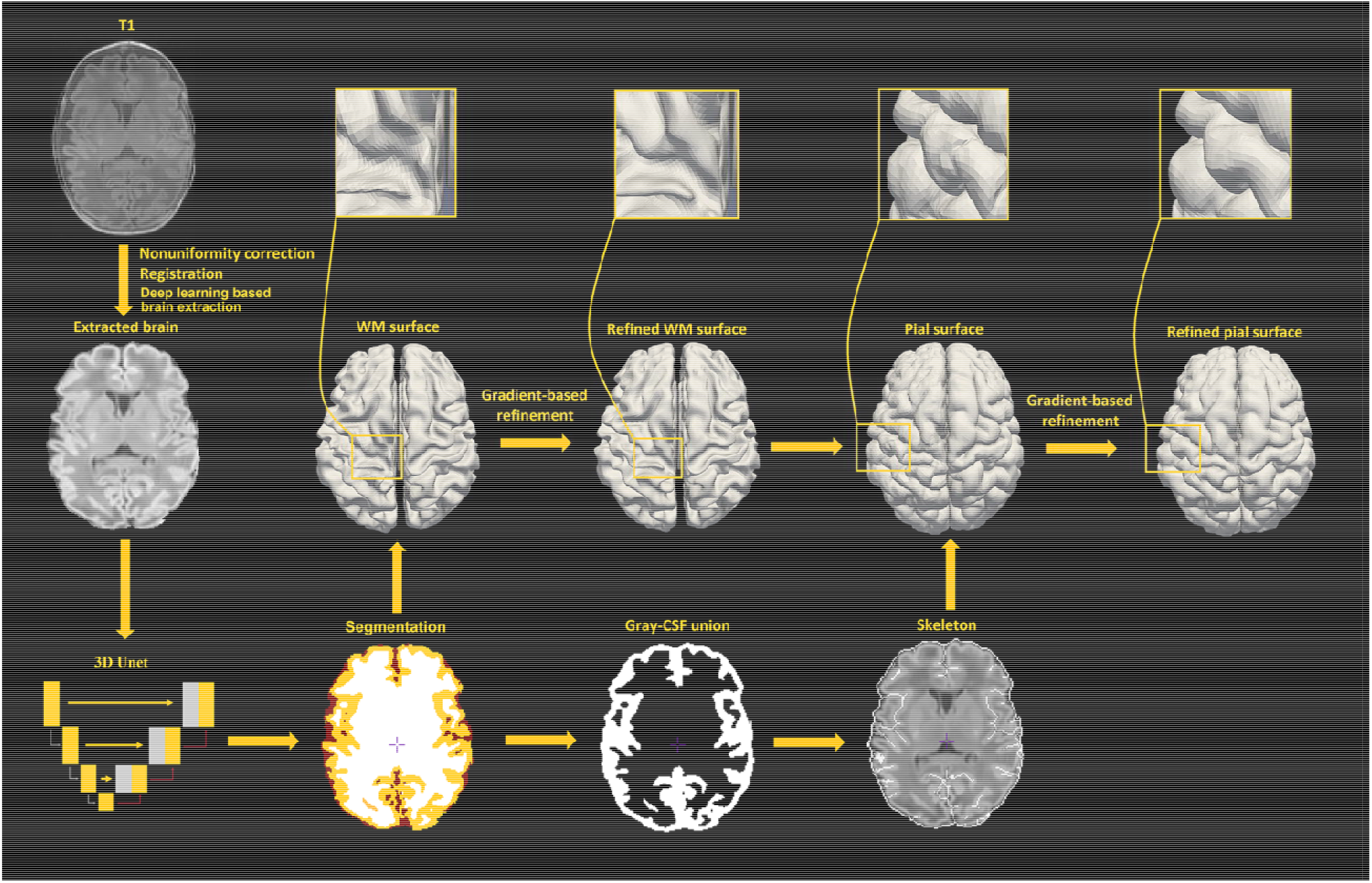
An illustration of pial surface reconstruction pipeline steps in this study.

### Brain extraction

Brain extraction is essential for T1w based neonatal surface reconstruction. Due to the similar intensity between WM and CSF, a liberal brain mask may include more CSF, which would increase the risk of mis-segmenting WM and CSF. In this study, a deep convolutional neural network-based approach, i.e. 3D-Unet (Çiçek et al., 2016), is used for brain extraction. 3D-Unet has been demonstrated to perform better than most of the existing methods in brain-extraction (Hwang et al., 2019). In our study, a 3D-Unet consists of three contracting encoder layers to analyze the whole image and three successive expanding decoder layers to produce a full-resolution segmentation (Ronneberger et al., 2015). Each layer contains two 3 × 3 × 3 convolutions each followed by a rectified linear unit (ReLu), and then a 2 × 2 × 2 max pooling with strides of two in each dimension. The 3D-Unet was trained on 36 × 36 × 36 voxels image patches fragmented from template MRI images.

We include 24 templates (MRI + manually segmented brain masks) from all age periods (24-45 weeks of PMA) to capture cortical morphology that dramatically changes during perinatal development. The templates used for the training of the segmentation algorithm were selected from all three datasets, and evenly selected across various postmenstrual age (PMA) groups (see details of template selection in Fig. 3). Manual segmentation of brain tissue is also included in the 24 templates, which are utilized as training templates for brain tissue segmentations, as discussed in the following section (see illustration of all 24 manual segmentations overlaid on MRI images in Fig. S2).

**Fig 3.**
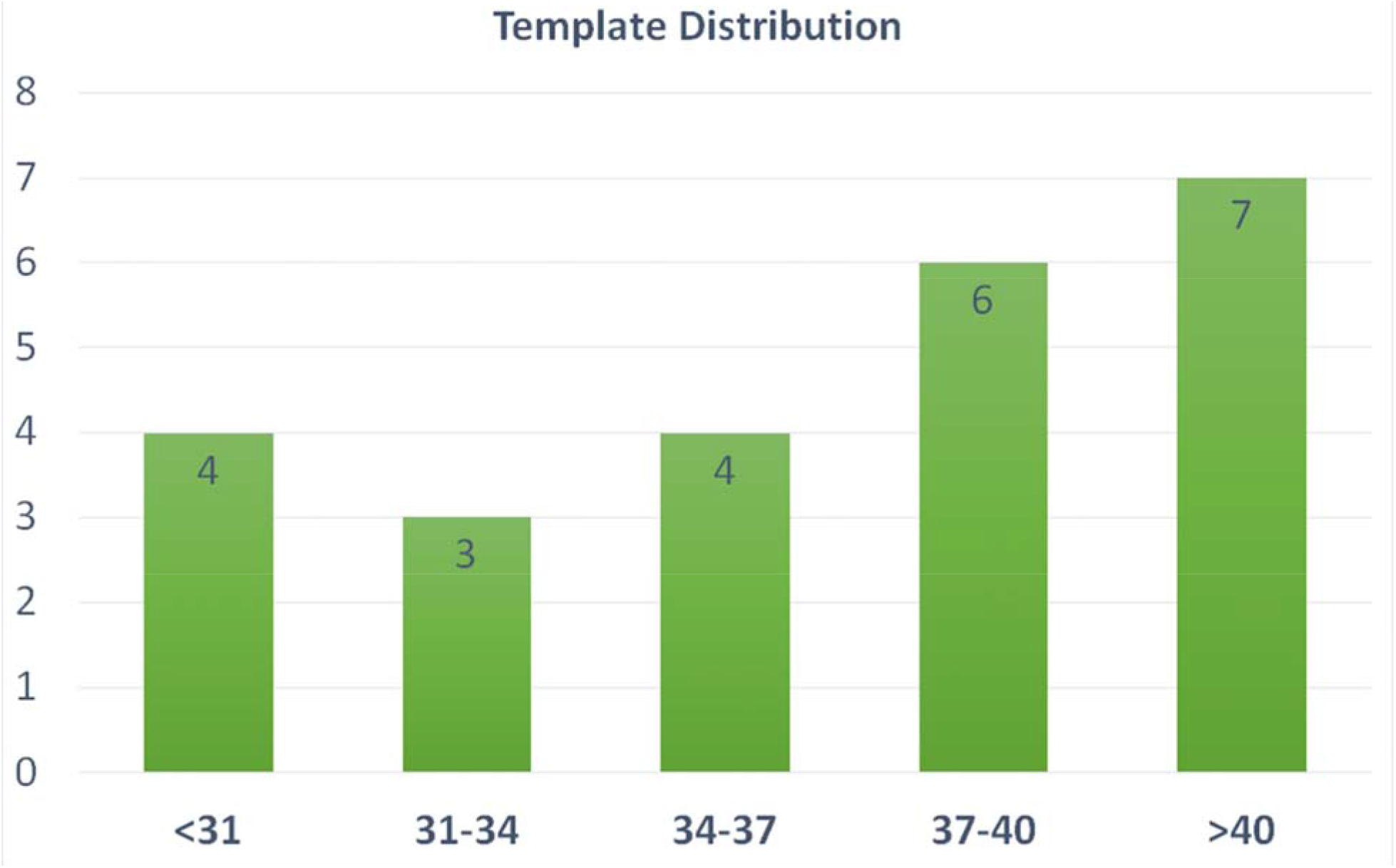
Age distribution of the templates selected in our pipeline. These templates are utilized in both brain extraction and tissue segmentation. X-axis label represent the PMA weeks.

### Tissue segmentation

Brain tissues are segmented into WM, GM, and CSF. Our previous work (Kim et al., 2016;Liu et al., 2019), despite successfully solving the issue of large morphological alterations, did not fully address the mis-segmentation between WM and CSF due to the confounding intensities between WM and CSF in neonatal MRI (Li et al., 2019) for neonatal brain (Fig. 4). To solve this, a non-local 3D-Unet (Wang et al., 2020b) is applied for tissue segmentation in our pipeline. Conventional 3D-Unet (like the one used in brain extraction) applies small kernels in convolution operators to scan inputs and extracts local information. As complex biomedical image segmentation usually benefits from a wide range of contextual information, one has to develop new operators that perform the global information aggregation to recover more accurate details. Non-local 3D-Unet was designed by adding a self-attention block, which computes outputs at one position by attending to every position of the input, to both the encoder layer and the decoder layer to aggregate/capture the contextual information. Non-local 3D-Unet has been demonstrated to outperform most state-of-the-art methods in neonatal tissue segmentation (Wang et al., 2020b).

**Fig 4.**
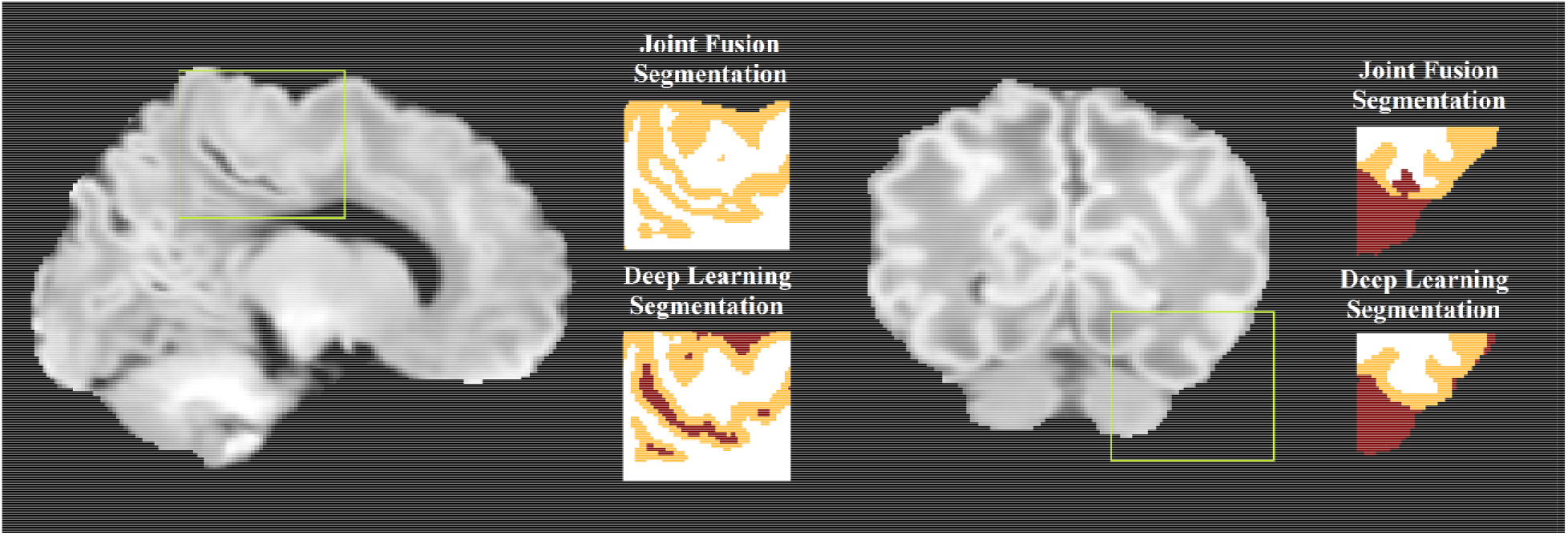
An illustration of mis-segmentation using joint fusion-based multi-atlas tissue segmentation. In the areas with dark WM, the surrounding CSF voxels are often misclassified into WM (left) or the dark WM voxels are misclassified into CSF (right), which becomes corrected using the proposed deep learning approach.

In our study, the non-local 3D-Unet consists of three encoder layers and three decoder layers. Each layer contains two 3 × 3 × 3 convolutions followed by a 2 × 2 × 2 max pooling with strides of two in each dimension, and an aggregated down/up-sampling self-attention block. The non-local 3D-Unet was trained on 36 × 36 ×36 voxel image patches fragmented from template MRI images. Fig. 4 illustrates that compared to the multi-atlas joint-fusion tissue segmentation that was applied in our previous work, mis-segmentation between WM and CSF is largely addressed by the non-local 3D-Unet.

### WM surface reconstruction

The WM/GM interface is extracted using the surface extraction tools employed in CIVET-2.1.0, with extensions to neonate brains. In the first stage, a new marching-cube approach is utilized, where the number of surface-meshes are adapted to fit the individualized brain size and morphometry. This could facilitate the parameter selection which is needed in the parametric deformable models (MacDonald et al., 2000;Kim et al., 2016), and avoid underfitting due to suboptimal parameters. Spherical topology of the surface is guaranteed by collapsing, in voxel space, an outer ellipsoid enclosing the WM mask onto this mask, while preserving the spherical topology of the ellipsoid. Automated mesh adaptation tools (in-house software within CIVET-2.1.0) are then used to coarsen the initial marching-cubes surface to a manageable size before inflating it to a sphere. Specifically, topologically valid vertices are removed by merging two vertices and one edge, and connections between new vertices and other vertices are setup to form new triangles. Then, the brain mesh is geometrically smoothed iteratively until a convex blob is obtained by an inflation algorithm, and the coordinates are normalized to the unit sphere. The inflated sphere serves as a mapping between the icosahedral sphere to the inflated sphere for resampling the brain surface like an icosahedral sphere at 81,920 polygons, as required by our in-house surface-based registration tools (Robbins et al., 2004). The resampled surface is then fitted to the T1w image. During the mesh deformation, a self-proximity term is also added to prevent the polygon intersection (Kim et al., 2005). In the second stage, the resultant boundaries are deformed again to refine the fitting using an image gradient term (see details in Gradient-based surface refinement). Examples of such improvements with the gradient feature-based refinement are shown in Fig. 5. This surface is registered to an age specific template using a spherical registration method and resampled into 81,920 meshes by the icosahedron resampling (Kim et al., 2016), more details can be found in section *Surface registration to age-specific templates*.

**Fig 5.**
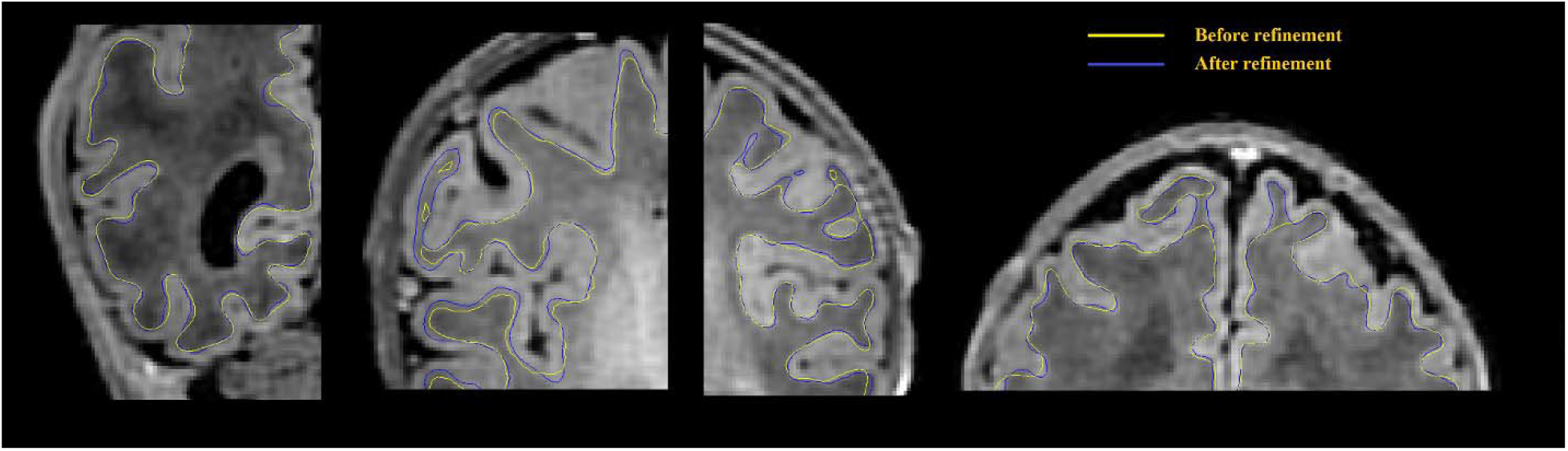
WM surface is significantly enhanced (visually closer to true GM-WM boundary) after gradient-based refinement. Yellow curves represent the WM surface before refinement, and blue curves represent the WM surface after refinement.

### CSF Skeletonization

The purpose of the skeletonization is to identify the hypothetical location of CSF within the sulcal bank and fundus to permit deep surface penetration of the pial surface in sulci. Most previous methods have attempted to detect the combination of CSF voxels and CSF partial volumes buried in the sulcus, and skeletonize them (Han et al., 2004;Kim et al., 2005;Lerch and Evans, 2005;Im et al., 2006;Lee et al., 2006;Xue et al., 2007). However, given that CSF voxels or CSF partial volumes can hardly be seen within a number of sulci on neonatal MRI due to small brain size, indistinguishable sulcal banks, and limited image resolution, the aforementioned approaches failed to reliably capture sulcal skeletons. Therefore, we choose to identify the medial surface of the union of GM and CSF as an alternative to CSF skeletons. The skeleton may not necessarily locate in the middle part of the GM-CSF union, therefore we use a second model to refine the initial pial surface, which is deformed to the skeleton, based on image intensities alone. If there are intensity gradient in unobserved CSF in the deep basin area, the skeleton is not a necessity to detect the CSF, since the difference of gradient and surface refinement would ultimately be used to detect the CSF. However, the skeleton has a strong effect to detect CSF in non-gradient or invisible CSF of the sulcal basin area, where the GM is hypothesized to have the same thickness in two sulcal banks.

The main steps involved in skeletonization are 1) Finding external and internal boundaries. We define the external boundary of the GM-CSF union as the hull of the brain mask and the internal border as the GM/WM boundary that is obtained from the white surface. Voxels located at the external boundary are first selected as part of the skeleton that represents the gyral interface. We then skeletonize the remaining part of the GM-CSF union by thinning the aggregation of voxels in the GM-CSF union, leading to the generation of medial surfaces of sulci. 2) Initialization of catchment basins for watershed algorithms. The latter skeletonization is conducted using homotopic erosion that preserves the initial topology, with a watershed algorithm embedded in the erosion process (Riviere et al., 2002). The first step in the watershed algorithm is to identify initial seed voxels for catchment basins to represent potential sulcal *fundi*. These seed voxels are determined by thresholding the Gaussian curvature for each voxel on the boundary of WM and GM-CSF union. 3) Recursive erosion of the GM-CSF union. Then, water rises from the catchment basins, which is driven by the mean curvature. Voxels on the edge of the GM-CSF union are removed if their mean curvatures are lower than a given threshold. This erosion is processed iteratively. A skeleton is generated when, by recursive erosions, the water reached ridges which correspond to the medial localization of cortical folds. 4) Pruning of the remaining voxels. Due to the excessive detection of initial seeds in the watershed algorithm, redundant skeleton branches may be created (Shen et al., 2011). Therefore, an additional step of pruning is applied to remove small non-significant secondary branches and construct a more reasonable skeleton of sulcation. Our pruning process is performed by classifying the remaining voxels as deletable points and undeletable points depending on their topological characterizations, i.e. how the voxels connect to their neighbors within the remaining voxels. Specifically, each remaining voxel is labeled as one of nine categories: *interior* voxel, *isolated* voxel, *border* voxel, *curve* voxel, *curves* voxel, *surface* voxel, *surface-curve* voxel, *surfaces* voxel and *surfaces-curve* voxel. The definition of the nine classes can be found in (Malandain et al., 1993;Mangin et al., 1995). The *surface* voxel, *surface-curve* voxel, *surfaces* voxel and *surfaces-curve* voxel are undeletable voxels; the *isolated* voxels are deletable voxels. For the all other categories, users are able to select which categories are deletable voxels (see *Parameters selection* section). The deletable voxels are iteratively removed until all the remaining voxels are undeletable voxels. Fig. 6 shows an illustration of skeletonization of the union of GM and CSF voxels.

**Fig 6.**
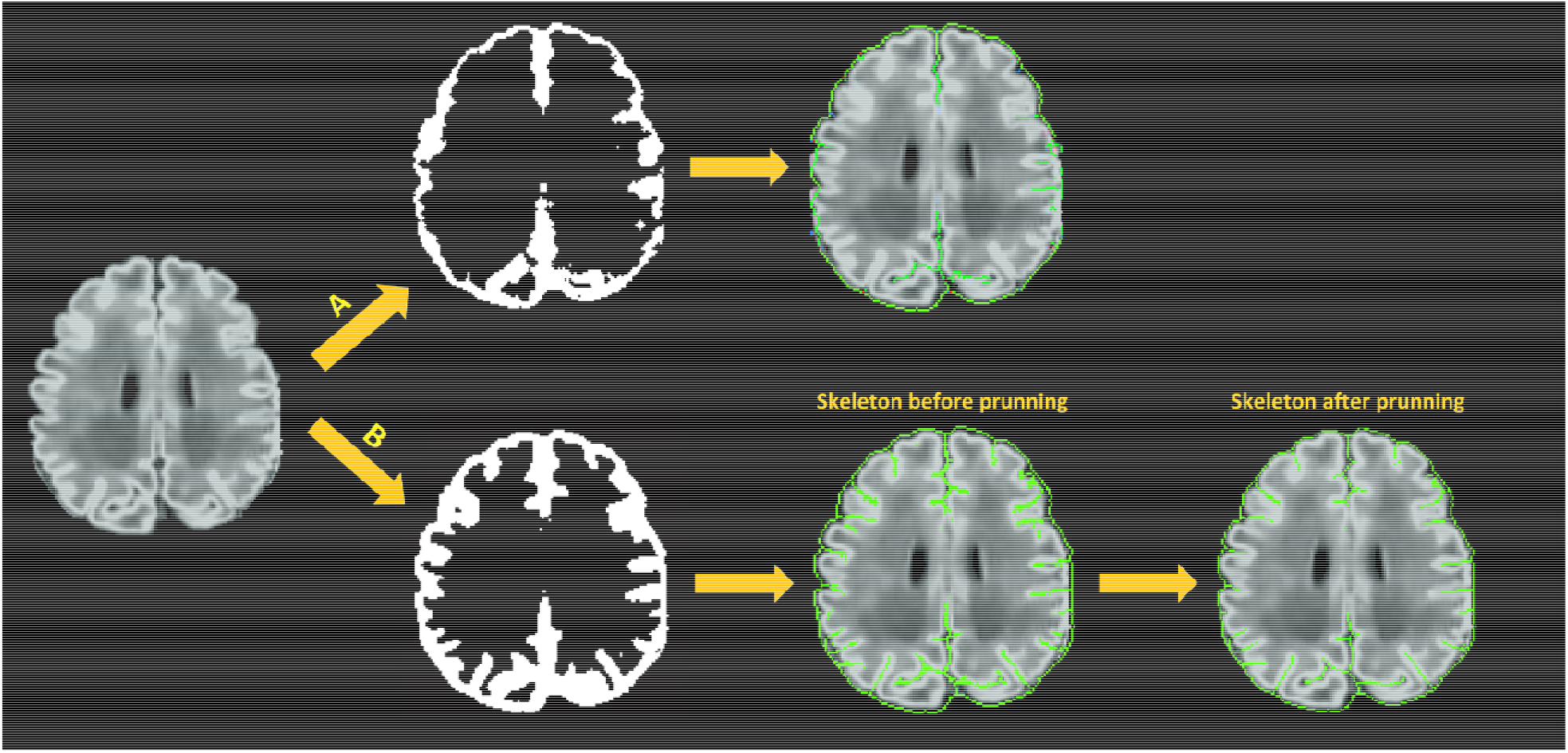
Skeleton generation within sulci with no CSF-PV; A) skeletonization of CSF voxels (CSF-PV>0%) only; B) skeletonization of the union of GM and CSF.

**Fig 7.**
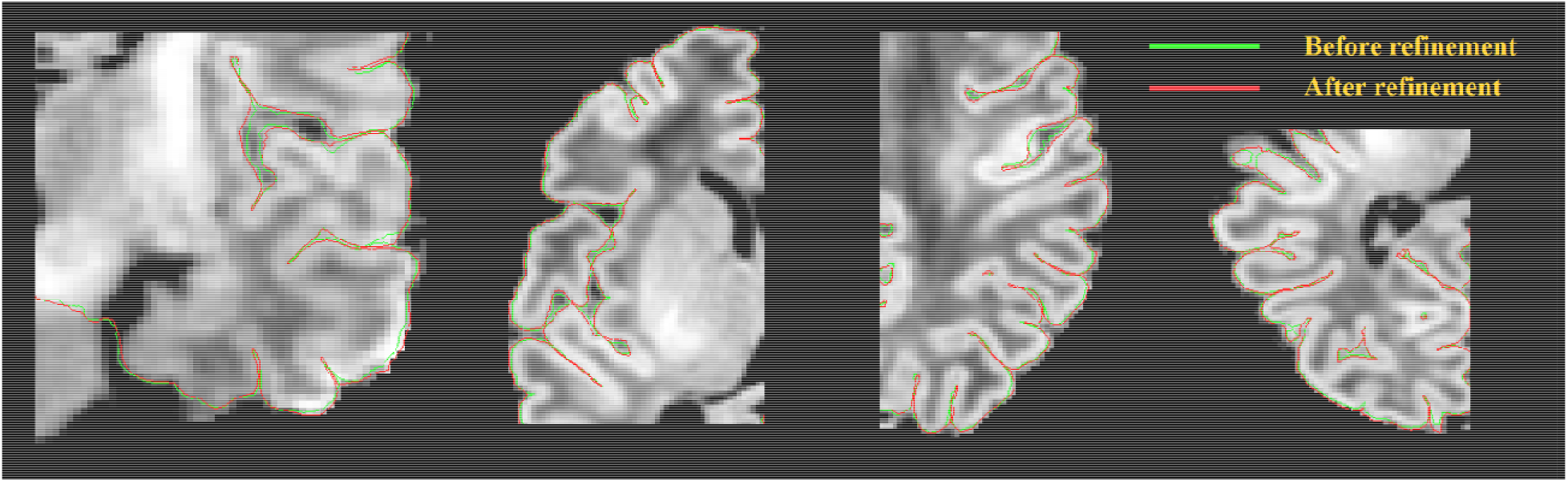
Pial surface significantly enhanced (visually closer to true GM-CSF boundary) after a gradient-based refinement. Green curves represent the pial surface before refinement, and red curves represent the pial surface after refinement.

**Fig 8.**
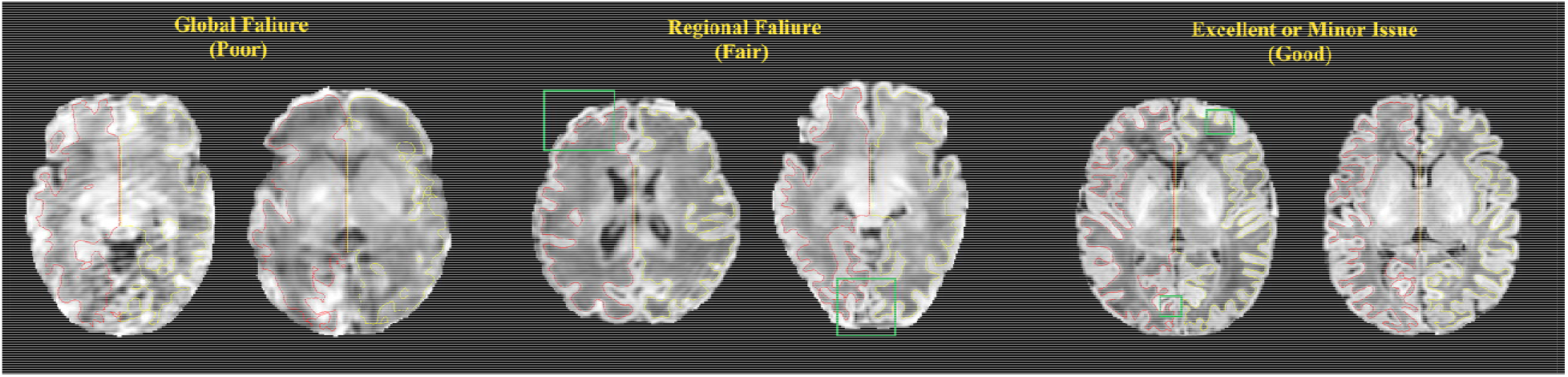
Examples of surfaces generated with poor, fair, and good qualities. These criteria match those used by our experts to manually score the quality of the surfaces.

### Pial surface reconstruction

The generation of pial surfaces consists of two stages: First, the initial pial surface is obtained by iteratively expanding the WM surface towards the skeleton of GM/CSF union. The expansion path to this boundary is defined using Laplacian fields generated from the WM surface to the skeleton (Kim et al., 2005). The path defined by the Laplacian fields provides the one-to-one projection/mapping of all the vertices on the WM surface onto the deformed surface. Defining the point correspondence between the two surfaces in this manner helps avoid the topological errors (Kim et al., 2005). On the other hand, the skeleton generated from the GM/CSF union is not a precise representation but rather a hypothetical delineation of the true location. In regions where CSF voxels or CSF partial volumes are observed within the sulcal bank, pial surface reconstructions based solely on the skeleton of GM/CSF union may not fit these CSF voxels. In the second stage, the pial surface is thus iteratively fine-tuned to fit the adjacent, more realistic GM-CSF boundary using gradient information derived from intensities of voxels proximal to the surface for each current iteration. To this end, we generate a gradient profile along the direction normal to the surface and converged the surface fitting to a position of the corrected local maximum gradient (see more details in *Gradient-based surface refinement*).

### Gradient-based Surface Refinement

Potential inaccuracy of the pial surface reconstruction might be generated when based solely on the skeleton of the GM/CSF union. These occurrences make sulcal CSF skeletonization more difficult, leading to a defective surface deformation toward the true cortical boundary. Therefore, we adapt, from CIVET-2.1.0, a supplementary model to improve the surface deformation, which is based solely on the image intensity gradients and is applied to both WM and pial surface fittings.

For a more accurate surface deformation, we strategically compute a directional gradient by analyzing the one-dimensional (1D) intensity profile along the radial lines perpendicular to th surface, at each vertex. This directional gradient is sampled equally in spatial points by a *search step* (e.g., 0.5mm) along the surface-normal rays within a certain distance range, defined a search *distance*, both inward and outward (e.g. −5mm to +5mm). The intensity values are blurred along each radial line to remove small local extrema. Raying from the center located on a given surface vertex, we perform a search process to identify a suitable edge (maximum gradient) among all the sample points within the defined search distance. For each vertex, the intensity at the sample point where the maximum gradient among all the sample points occurs is recorded. In this manner, we could map such intensities across all vertices. By doing so, the local target intensity, which is used to compute the maximum intensity gradient, is allowed to vary over the entire surface. Then, surface-based diffusion blurring is applied to this intensity map by incorporating information from neighboring vertices. This blurring procedure is critical because the extracted gradient information along the surface-normal profile is occasionally erroneous due to motions, image artifacts, bias field, blood vessels, flawed surface normal, and a relatively small range of the search distance. This blurring process attenuates erroneous gradients or outliers, thereby leading to more robust convergence. Finally, the surface mesh is deformed using 1) forces of attraction to locations displaying maximum blurred gradient values; and 2) constraint of the vertex-wise spatial smoothness. Our proposed gradient-based deformation follows a multi-resolution approach to efficiently fit the initial surface to the location displaying the maximum gradient, while avoiding being trapped in local maxima. Parameters related to the multi-resolution approach are described in *Parameters selection* in supplementary materials.

### Surface registration to age-specific templates

We register individual surfaces to the template using *SURFTRACC* (Robbins et al., 2004). This tool first transposes individual sulcal depth maps into icosahedral spheres. It then iteratively searches for local optimal vertex correspondence between an individual and a template sphere based on a matching of a given feature (in this case, depth potential function (Boucher et al., 2009)). Mapping the deformed meshes back to the original cortical surface coordinates allows a registration of the individual to the template surface. To address for cortical folding changes dramatically during perinatal development, we previously constructed age-specific surface templates representing four age ranges: 26-30, 31-33, 34-36, and 37-40 weeks PMA. Details are found in our previous study (Kim et al., 2016). For inter-subject analyses and group comparisons, any given subject is registered to its corresponding age template and then is ultimately transformed to the oldest template space by concatenating the sequence of transformations between age-specific templates. It is noted that neonatal brains with pathology or an injury like intraventricular hemorrhage, white matter injury or ventriculomegaly may under-develop or deviate from the normal development (Kim et al., 2020). Use of their postmenstrual age (PMA) in finding their age-matched template may thus lead to a suboptimal registration result. Also, there may be cases where their PMA are not provided. To overcome this, we first register the given individual surface separately to each of all the four age-specific templates. We then identify the best-matched template based upon the similarity (using Pearson’s correlation) of each template’s sulcal depth pattern with the registered individual surface. To evaluate this, we compare the age of the template chosen by the proposed method with that matched with the subject’s postmenstrual age. We perform this comparison in preterm neonates with injuries and those without.

## Results

### Surface quality assessment

The qualities of WM and pial surfaces were assessed by visually scoring the fitting accuracy of the reconstructed surfaces. The scoring was done by assigning an overall score from 1 to 3 to each of the individual pial surfaces. Score 1 indicated poor quality, in which the contour substantially deviated from the cortical boundary, and the deviation was distributed globally. This occurs mostly due to severe motion artifacts. Score 2 indicated cases where the contour was close to the cortical boundary, but contained several local flaws. This occurred in cases with minor motion artifacts, and failed segmentation in the full depth of narrow sulci fundi. Score 3 indicated a contour with accurate tracing of the cortical boundary, or with only focal and slight deviation, which was difficult to find without careful review. Note that we considered surfaces with minor issues as good quality (score 3), which may have resulted in more ‘good’ cases in our evaluation than previous studies (Makropoulos et al., 2018b). Two independent raters performed the quality assessment (QA) for all three sets (J. S. & N. T.) respectively.

To quantitatively perform the surface QA, we further conduct a comprehensive landmark-based analysis of accuracy for outer cortical surfaces. An independent rater (J.K.) visually picked a number of landmark points located on the GM/CSF interface. These landmarks were considered as samples representing the “ground truth” surface boundary. The Euclidian distance from each landmark to either the pial surface or the intermediate pial surface (generated at stage 1 where no gradient information was used) was computed. Due to shallow cortical folding, fewer landmarks are selected in younger preterm neonates such that we select 30 landmarks in the group of 26-31 weeks PMA, 50 in 32-34 weeks, 100 in 35-36 weeks, and 100 in 37-45 weeks. The landmarks were selected randomly, but evenly distributed across cortical lobes (i.e. include but not limit to frontal, parietal, temporal, and occipital lobes) and approximately evenly distributed across all the sulci (i.e. mainly distributed in several main sulci like central sulcus, lateral sulcus, cingulate sulcus, and parietal occipital sulcus) that showed a sufficient depth and an eminent shape relative to secondary branches. Due to a large amount of time required for this manual procedure, 20 neonates (randomly selected with PMA evenly distributed between the aforementioned 4 age groups) were selected from the three datasets.

#### Qualitative evaluation results

Fig. 9 presents the results of the surface QA for three datasets. WM surfaces produced by the proposed method were rated as good/accurate (score 3) in 83.6% of scans (averaged by two raters). Pial surfaces were rated as accurate in 82.1% overall. For the UNC dataset, WM surface yielded 92.1%, and pial surface 89.5%. For the dHCP dataset, WM surface yields 92.2%, and pial surface 89.2%. These results indicated no visual mistakes on the majority of the constructed surfaces. The surfaces rated as fair (average across three datasets: 7.8 % for WM surfaces; 9.7% for pial surfaces) showed regional flaws but were similar to the true cortical boundary (score 2). Only 2.8% of the WM surfaces and 3.4% of the pial surfaces (average across three datasets) displayed poor quality (score □ = □1).

**Fig 9.**
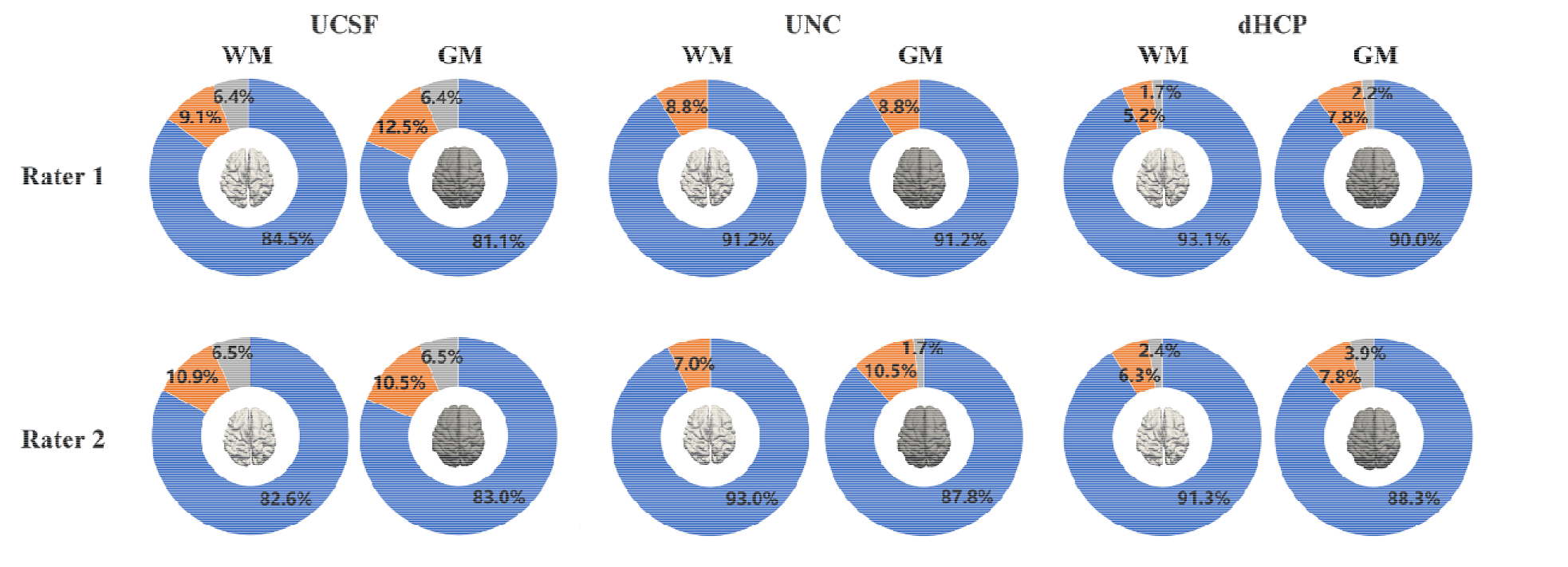
The results of the WM and pial surface QA for all three datasets. Blue represents the proportion of individual surfaces with score 1 (good quality), orange score 2 (fair quality with regional defects) and gray score 3 (poor quality). WM surfaces produced by the proposed method were rated as accurate in 89.2% of scans overall (averaged across three datasets by two raters). Pial surfaces produced by the proposed method were rated as accurate in 86.9% overall.

Inter-rater comparison was also performed to determine how many surfaces are consistently given the same score between the two raters”. Results indicated that among 843 scans (acros three datasets), 116 WM surfaces (13.8%) were rated differently, and 141 pial surfaces (16.7%) were rated differently, which suggest the surface QA was relatively consistent across raters.

#### Quantitative evaluation results

Analyzing the 20 randomly selected subjects, the landmark-based displacements for the intermediate pial surfaces and refined surfaces are shown in Fig. S3. Results exhibited that the mean displacement for intermediate surfaces (mean sd = 0. 0.049mm) significantly decreases when they were refined using the gradient information (0. 0.035mm; p<0.0001, paired t-test).

### Selection of age-specific templates for the surface registration: using sulcal depth similarity vs. postmenstrual age

We evaluated here whether the template yielding the highest correlation of sulcal depth measurement with the given individual surface to register is concordant with the template chosen using the PMA of the given individual data. To this end, we constructed a confusion matrix to map their overlap (Fig S4). Results indicated that overall 82.0% of our data showed overlap between the two methods. Most of the mis-matched templates (15.6%) were found in the nearest neighboring age groups. Among the rest of the mis-matched templates (2.4%), the majority were those with brain injury (75%), whose matched templates were identified to be younger than their PMA. The overlap was lower for the youngest and the oldest babies (65-69%) than other age groups (90-94%).

### Surface reconstruction in dHCP measurements

To directly compare between the quality of surfaces reconstructed by our pipeline and by th dHCP pipeline, we randomly selected 12 images from the dHCP data set and a trained expert (M. L.) drew landmark points on the white matter and pial surfaces based on their T1w MRI. On average, 80 landmark points were placed on both of these surfaces, in both hemispheres (see Fig. 10 for an example). We then computed the shortest distance between these points and our computed surfaces. Additionally, in the case of the dHCP dataset, we also computed distances between the manually drawn landmark points and the surfaces provided by the dHCP minimal processing pipeline. The mean and standard deviation of the absolute value of these measurements are included in Fig. 11. The white matter surfaces tended to be comparable between the two pipelines, while in the case of the pial surfaces, our pipeline yielded lower distance with the landmarks taken from the T1w image. That was, our pipeline showed significant different placements of the pial surface from the dHCP solutions which extracted cortical surfaces mainly on the T2w image (p<0.05).

**Fig 10.**
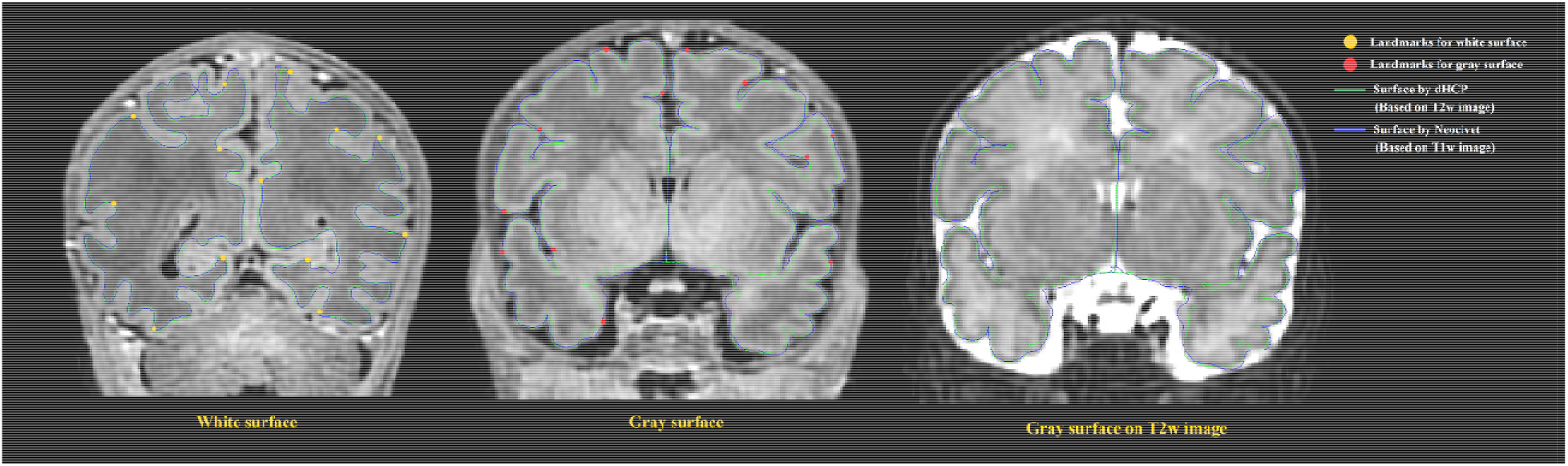
Examples of surface validation points (landmarks) placed on the WM/GM boundary and the GM/CSF boundary of a randomly selected T1 weighted image from the dHCP dataset. **Left:** Surface validation points from the WM surface are shown along with the dHCP and our WM surface reconstruction solutions; **Middle:** Validation points from the pial surface are shown along with the dHCP and our pial surface reconstruction solutions. **Right:** Pial surface were mapped to same MRI slice from the dHCP dataset as shown in middle panel but with T2 modality. Note that the dHCP surfaces are extracted on T2 weighted images that may estimate CSF partial volume and pial surface differently. The dHCP surfaces are indeed observed to fit better the pial boundaries found on T2 weighted images than those on T1 weighted.

**Fig 11.**
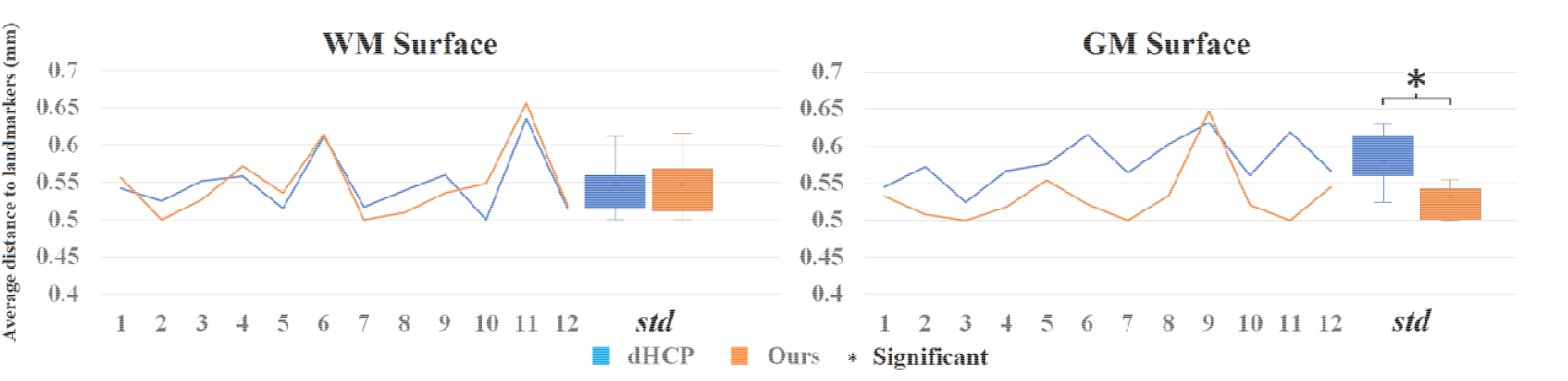
Mean distance (mm) to the landmark from the dHCP surface (blue) and our surface (orange) for 12 random selected subjects from dHCP dataset. Note that the dHCP surfaces are extracted on T2 weighted images that may estimate CSF partial volume and pial surface differently.

Note that the surfaces extracted on T2 weighted images that may estimate CSF partial volume and pial surface differently compared to T1 weighted images. Fig. 10 right panel shows that the same slice as the one showed in Fig.10 (middle) but with T2w modality.

### Cortical thickness measurements

To evaluate the ability of our pial surface reconstruction in characterizing the hypothesized developmental trajectory, we measured cortical thickness by the shortest distance between pair of vertices on the WM and pial surfaces. The vertex-wise thicknesses were calculated in the native MRI space where the scan was acquired.

We mapped the individualized cortical thickness trajectory with age for both UCSF and dHCP datasets in whole brain and 6 anatomical lobes. We observed a very smooth overlap between the two datasets and the trajectory of cortical thickness changes along with aging (Fig. 12B). Across the two datasets, our pipeline identified increases in cortical thickness over time in almost all cortical regions (overall cortical thickness: +0.041 mm/week; p < 0.0001, t = 4.99; Fig. 12A). A vertex-based mapping showed that regions involved in the cingulate cortex display the most rapid growth among all brain regions whereas brain regions in the inferior temporal lobe, exhibited the slowest cortical thickness growth. Fig. 13 presents cortical thickness maps in 4 cases representing each of the PMA age group.

**Fig 12.**
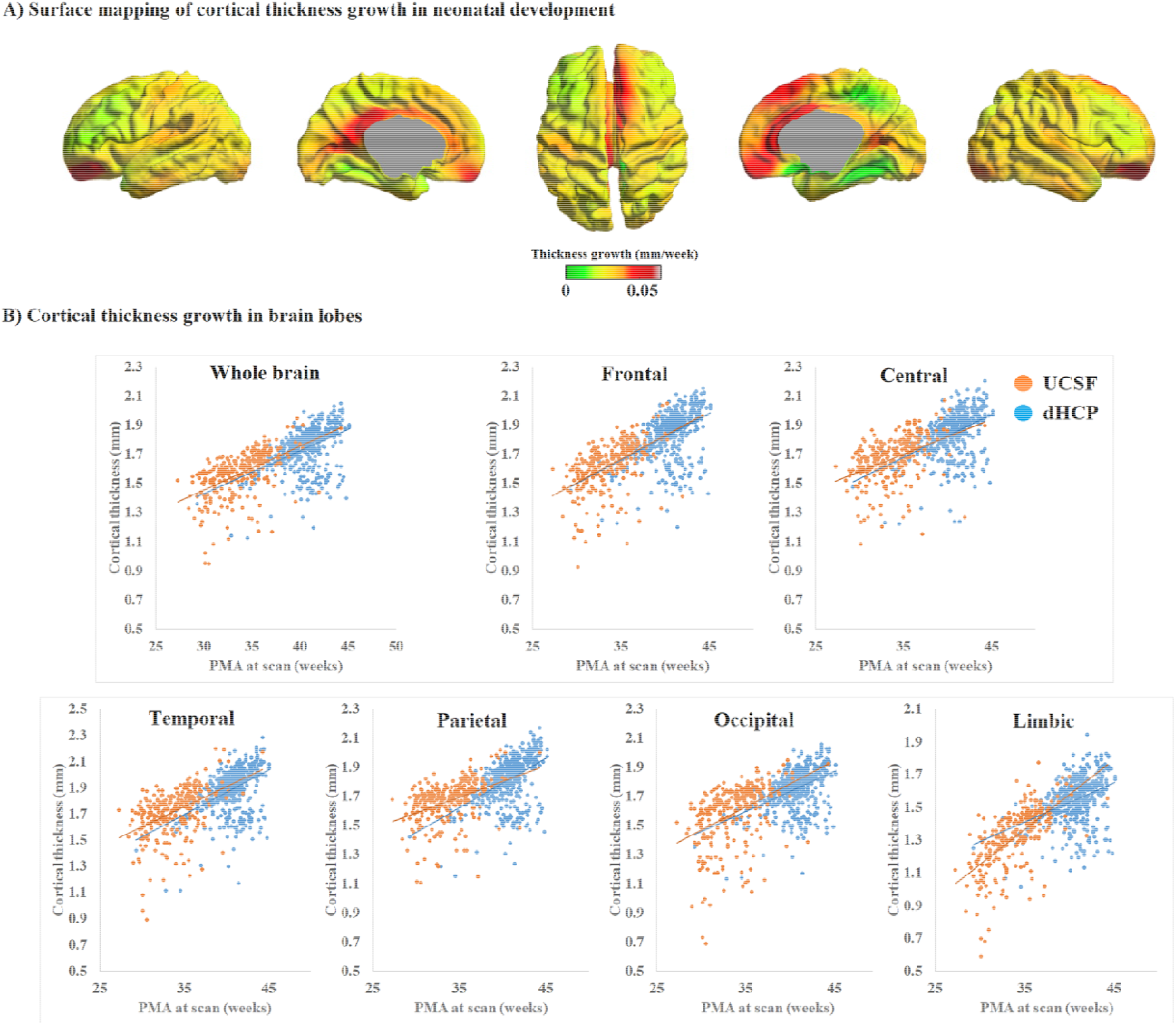
A) Mean cortical thickness significantly and positively correlates with PMA at scan (*p* < 0.001). Cortical thickness grows the most rapidly in brain regions near the cingulate cortex but the slowest in brain regions near the inferior temporal cortex. B) Trajectory of cortical thickness development in whole brain and 6 anatomical lobes, for both the UCSF and dHCP datasets.

**Fig 13.**
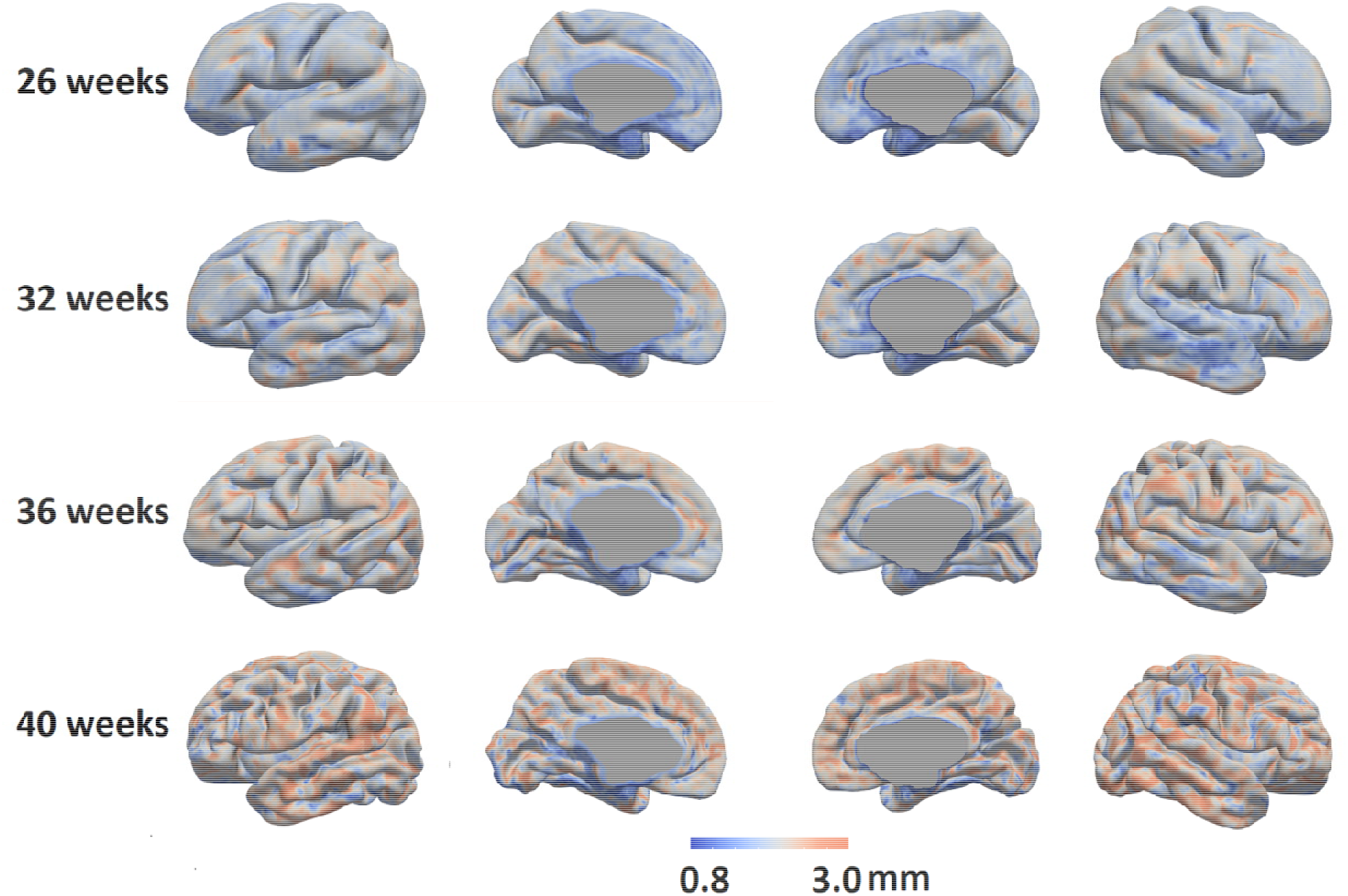
Cortical thickness maps in 4 individual cases representing 4 different PMA groups were shown (a: 26 weeks, b: 32 weeks; c: 36 weeks; and d: 40 weeks.).

### Comparison with literature

Cortical thickness measurements in the neonatal population have been reported in past literature and consisted of varying values, ranging between 1 and 2.2 mm (Xue et al., 2007;Moeskops et al., 2013;Li et al., 2015;Moeskops et al., 2015;Makropoulos et al., 2016;Geng et al., 2017;Makropoulos et al., 2018b). One reason for variability in measurements can be due to differences in image acquisition, image segmentation, and surface reconstruction methods. Fig. 14 maps the distribution of cortical thickness values measured from multiple studies in relation to their average PMA at scan.

**Fig 14:**
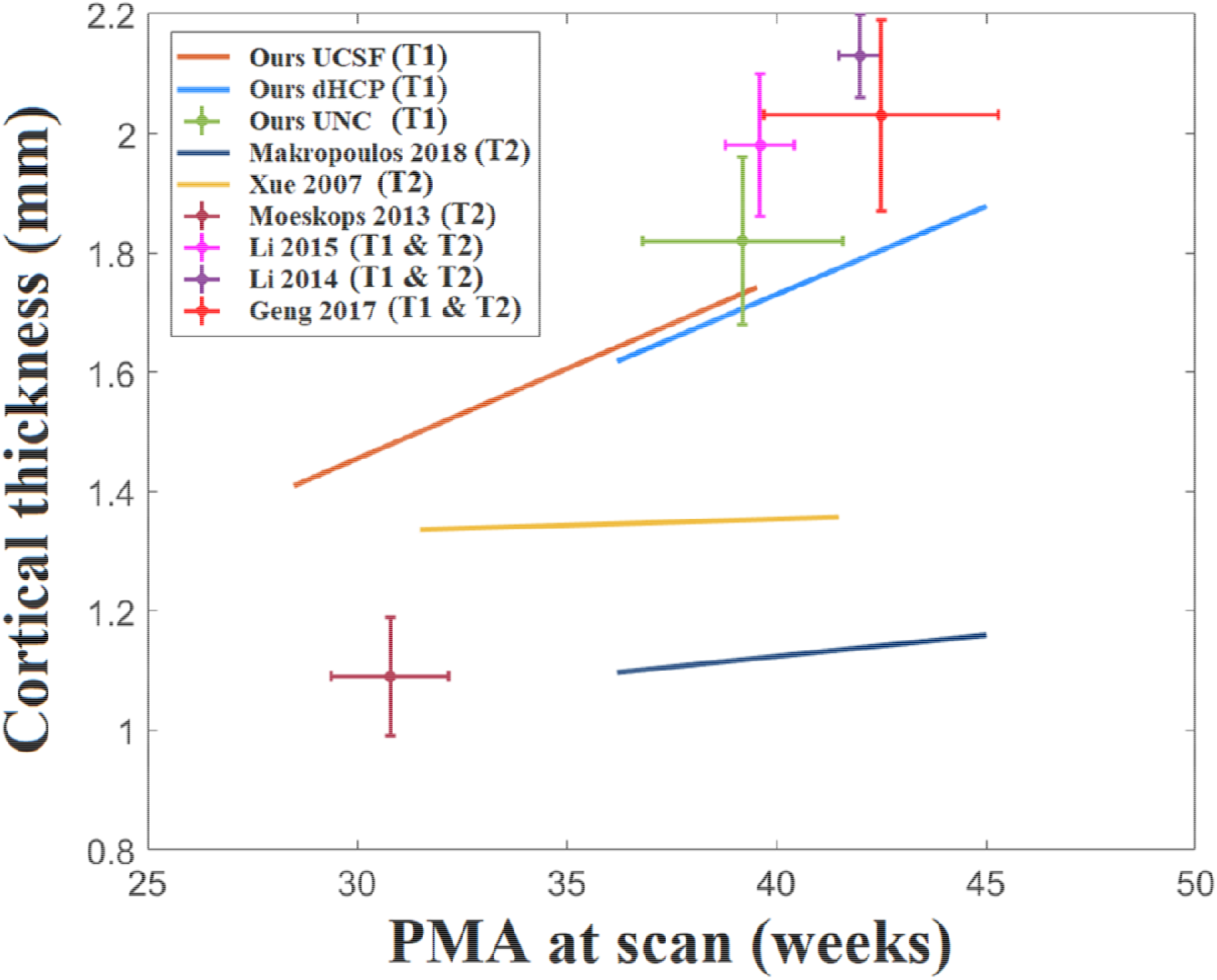
A statistic of neonatal cortical thickness measured from different studies. We consider the age range of the samples in each study and the growth trajectory. However, we found that the age ranges for some of the studies are very small. For example, Moeskops’s study only tested 10 subjects and the standard deviation of the age range is only 1 week. For those studies with relatively short age ranges (SD < 1.5 weeks), we indicate them with crosshairs that represent the two-standard deviation of the age range (horizontal extent) and cortical thickness (vertical extent). On the other hand, regression lines are used to represent the cortical thickness growth for those studies with larger age ranges (SD > 1.5 weeks).

### Comparison to the NEOCIVET 1.0 pipeline

Our new framework extended upon the NEOCIVET 1.0 pipeline (Kim et al., 2016) which wa designed for neonatal WM surface (WM-GM interface) reconstruction and measurement of sulcal depths. All new features that are incorporated are summarized relative to the NEOCIVET pipeline in **Table 1**.

**Table 1.**
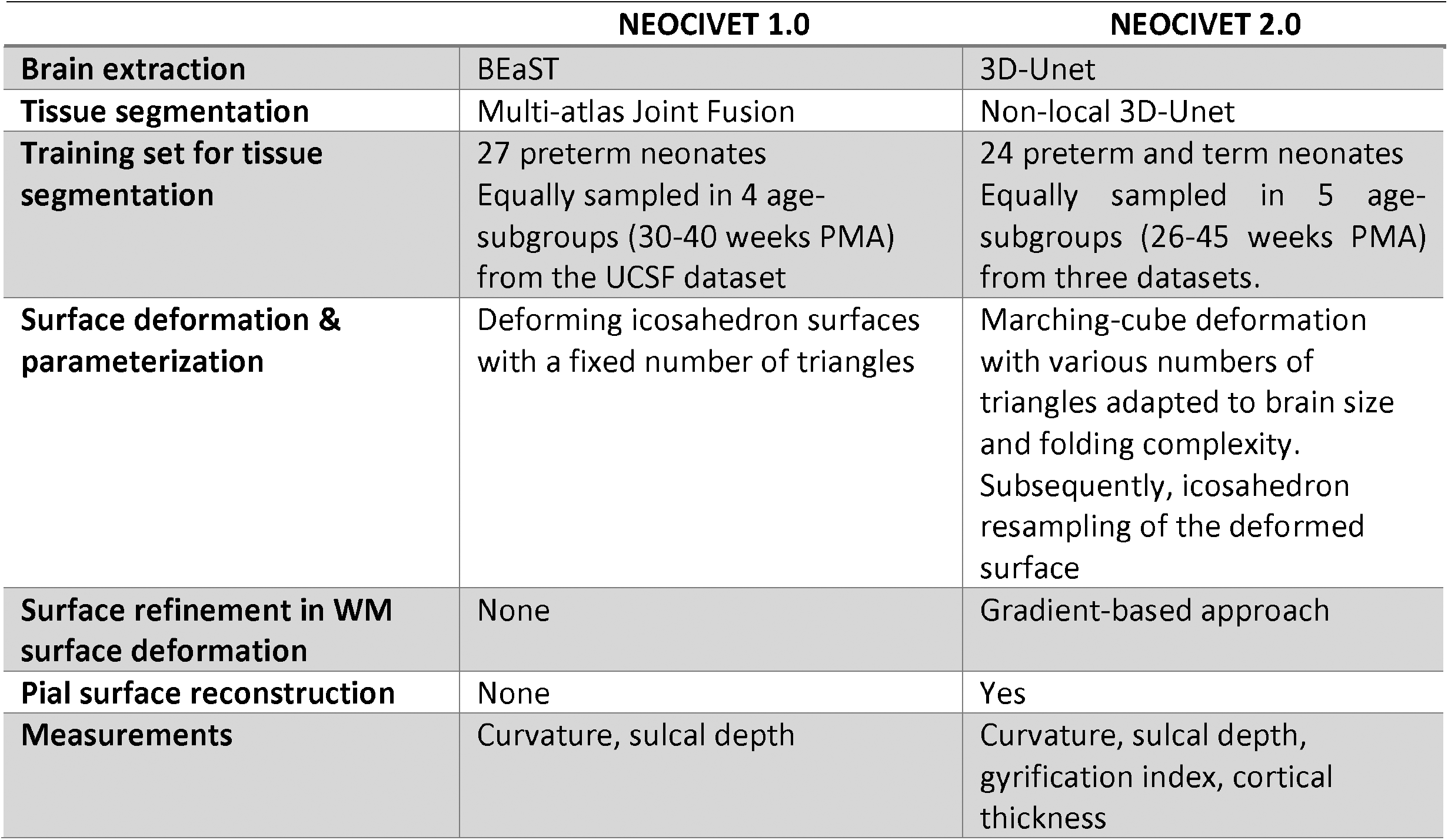
Comparison between NEOCIVET 1.0 and NEOCIVET 2.0 pipeline

| NEOCIVET 1.0 | NEOCIVET 2.0 |
| --- | --- |

|  |  |  |
| --- | --- | --- |
| **Brain extraction** | BEaST | 3D-Unet |
| **Tissue segmentation** | Multi-atlas Joint Fusion | Non-local 3D-Unet |
| **Training set for tissue segmentation** | 27 preterm neonates<br>Equally sampled in 4 age-subgroups (30-40 weeks PMA)<br>from the UCSF dataset | 24 preterm and term neonates<br>Equally sampled in 5 age-subgroups (26-45 weeks PMA)<br>from three datasets. |
| **Surface deformation & parameterization** | Deforming icosahedron surfaces with a fixed number of triangles | Marching-cube deformation with various numbers of triangles adapted to brain size and folding complexity.<br>Subsequently, icosahedron resampling of the deformed surface |
| **Surface refinement in WM surface deformation** | None | Gradient-based approach |
| **Pial surface reconstruction** | None | Yes |
| **Measurements** | Curvature, sulcal depth | Curvature, sulcal depth, gyrification index, cortical thickness |

Notably, to allow the new pipeline to better segment images scanned for old preterm or term neonates exhibiting large variability of cortical folding across individuals, we add more manually segmented templates from old preterm neonates and term neonates from the three datasets (PMA=40-45 weeks; n=7) in the library for tissue segmentation, which was not included in NEOCIVET 1.0. Finally, the new pipeline is able to extract the pial surface and measure cortical thickness, which are novel features added to the proposed pipeline.

We further compared the technical details between our pipeline and the dHCP pipeline, which can be found in supplementary materials.

## Discussion

In this study, we propose a new NEOCIVET pipeline (NEOCIVET 2.0) for automatically reconstructing neonatal WM and pial surfaces. Our new framework extends the NEOCIVET 1.0 (Kim et al., 2016), which was designed for neonatal WM surface (WM-GM interface) reconstruction. The NEOCIVET 2.0 pipeline primarily aims to strategically address important issues in neonatal MRI processing**: 1)** spatiotemporally changing tissue contrast**; 2)** significant partial volume effects which interfere with modeling sulcation in regions where detected CSF voxels are scarce and scattered. These issues could lead to defective pial surface reconstructions and inaccurate cortical thickness measurements if pipelines designed for adult brain MRIs, e.g., CIVET (MacDonald et al., 2000;Kim et al., 2005), FreeSurfer (Fischl, 2012), and Caret (Van Essen et al., 2001), were utilized. Specifically, to optimize neonatal pial surface reconstruction, we incorporate special features: 1) Construction of skeletons from the union of gray matter and CSF; and 2) Generation of pial surfaces by expanding the WM surface mesh towards the skeleton surface, a process refined using gradient information.

Furthermore, although NEOCIVET 1.0 successfully addressed several substantial issues/challenges of neonatal brain MRI processing (e.g., similar tissue intensity between WM and CSF, reduced tissue contrast between WM and GM, large within-tissue intensity variations, and regionally-heterogeneous image appearances that dynamically change along with neural development), our proposed framework included additional modules that further improved the quality of WM surface reconstruction. For instance, deep learning-based convolutional neural networks (Wang et al., 2020a) were introduced into the pipeline to render more accurate brain extraction and tissue segmentation, and to address the mis-segmentation between WM and CSF in neonatal T1w images. No existing pipelines that process neonatal brain MRI and perform cortical morphometry have not integrated deep learning neural networks for segmentation into their frameworks. We also incorporated a new marching-cubes approach that provides adaptive surface meshes according to each individual’s cortical folding complexity. The constructed surface is then further fine-tuned using gradient information extracted from voxel intensities.

Our qualitative assessment showed that our method can reconstruct most of the surfaces analyzed with reasonable quality. Our landmark-based quantitative assessment also suggests that the proposed gradient-based refinement method combined with skeleton fitting of CSF/GM union can more accurately position the pial surface to a location that is within a significantly small distance from the visually recognized “true” boundary. Similarly, the quantitative assessment also showed that NEOCIVET 2.0 fitted WM surfaces significantly closer to the “ground truth” in comparison to NEOCIVET 1.0 pipeline. Given such high accuracy as demonstrated in the evaluations, NEOCIVET 2.0 successfully characterizes and documents the developmental trajectory of cortical folding and thickness growth in preterm neonates. Furthermore, cross-validation using three independent datasets also suggests the generalizability of our proposed approach and the reproducibility of results using different datasets.

### Skeleton or non-Skeleton

Due to large partial volume effects in neonatal MRIs, CSF voxels within deep and narrow sulci typically cannot be captured with relatively large MR image resolution. Two main approaches, one with skeleton and the other without, were designed to circumvent this issue (Li et al., 2019) for neonatal pial surface reconstruction. However, both approaches encounter several issues. For example, a non-skeleton approach generates the pial surface based only on the edges of the identified CSF voxels and fails to consider the location of undetectable CSF. On the other hand, the skeleton approach may have a lower accuracy in properly pinpointing the correct positions of CSF (Osechinskiy and Kruggel, 2012), as it does not use edge information. As a result, both methods can lead to unreliable extraction of the sulcal boundaries.

Our approach combined skeletonization and edge information to improve the accuracy of the extraction of the pial surface. Regardless of invisible CSF voxels in sulci, skeletonization creates a barrier that can constrain the surface deformation. Furthermore, it provides a location where the GM-CSF boundary can be found within a very short distance. In this respect, our pipeline was able to more reliably search and refine true GM/CSF edges.

There have also been attempts to overcome the limitations associated with skeleton approaches. For example, a weighted geometric distance has been applied to the neonatal pial surface reconstruction (Xue et al., 2007;Li et al., 2012;Li et al., 2014) to solve this issue. This method, however, relies heavily on segmentation of CSF components, which can be severely limited in cases of large partial volume effects.

### Quality assessment of surface reconstruction

Landmark-based surface quality assessment (QA) showed that the displacement of our reconstructed WM and pial surfaces to the “true” WM and pial borders ranged between 0.5-0.65mm, which is quite small compared to the image resolution for scanning the UCSF and UNC datasets (0.6-1mm). This QA method, which was first designed to evaluate the adult brain surface reconstructions (Han et al., 2004;Tosun et al., 2006;Xue et al., 2007), was only performed in one neonatal/infant study in the past (Zöllei et al., 2020). Their pipeline was also evaluated on T1 images from the dHCP dataset. Landmark displacements for cortical surfaces measured in their study range between 1.17 to 1.19 mm for WM surface and 0.8 to 0.91 mm for pial surface respectively (Zöllei et al., 2020), which are much larger than that in the current study, implying the robustness of our proposed reconstruction method. Differences in the reported displacement values between our study and adult studies may stem from differences in image segmentation, image resolution, and brain size.

### Growth pattern in cortical thickness

Cortical thickness measurements (1.4mm at 28 weeks of PMA to 2.2mm at term equivalent) computed through our pipeline is quite similar to past studies (2.1 mm at term; e.g., Li et al., 2014) but different from others (1.1 mm at term) (Makropoulos et al., 2018b). This discrepancy may be explained by differences in thickness measurement methods, image modality (T1w vs. T2w), image segmentation, and surface reconstruction methods. Our results demonstrated a global increase of cortical thickness that positively correlated with PMA at scan. For the vertex-based ROI analysis, as in Figure 12B in the main text, we observed a very smooth overlap between UCSF and dHCP datasets, and the trajectory of cortical thickness changes along with aging are reasonably linear in both of the two datasets. In particular, brain regions corresponding to cingulate cortices grew the most rapidly. Similar results were reported in a past study which demonstrated that the cortical thickness is higher in the cingulate cortices and superior frontal cortex compared to other brain regions of term-born infants (Li et al., 2015), indicating that cortical thickness grows the most rapidly in these regions before birth. By contrast, brain regions in the temporal lobe grow the least rapidly. Cortical plate and subplate in these regions are found to grow more rapidly earlier in the 2^nd^ trimester of gestation (Vasung et al., 2016).

Other studies have identified alterations of cortical thickness and cognition in adolescents and adults born preterm, especially in the frontal cortices (Martinussen et al., 2005;Nagy et al., 2010;Nam et al., 2015). Thus, considering past studies combined with our results, regions near the cingulate cortices may be important biomarkers of cortical thickness changes in neurodevelopment, and these regions may be vulnerable to long-term alterations associated with prematurity or perinatal injuries that influence clinical outcome. Such alterations in cortical thickness development may be affected by both genetic and environmental influences (Lenroot et al., 2008) and have been associated with intelligence, executive functions, working memory, perceptual skills, and internalizing and externalizing behavior (Lohaugen et al., 2009;Skranes et al., 2012;Zubiaurre-Elorza et al., 2012).

Ultimately, cortical thickness trajectories have important clinical relevance and can be useful biomarkers that help to predict neurodevelopmental and cognitive outcomes. The new pipeline is capable of mapping dynamic cortical thickness changes throughout the third trimester of gestation, an early and critical phase of development, which can further support the early diagnosis of preterm neonates and, thereby, allow clinicians to determine when and what medical interventions may be appropriate.

### Limitations and future directions

We applied our pipeline to T1w images, while many attempts in neonatal brain segmentation and surface reconstruction were conducted on T2w images or T1w-T2w modalities combined (Shi et al., 2010;Wang et al., 2014;Makropoulos et al., 2016;Makropoulos et al., 2018b) due to the increased contrast of brain tissues in T2w images. It is important to optimize a method that considers T1w images, which are collected prevalently for various research aims. Our pipeline is the first proposed specifically for T1 images. Despite our new evaluation results where cortical thickness measurements vary depending on the image modality analyzed by each pipeline, it is difficult to conclude that whether T1 or T2 imaging is more ideal to construct cortical surfaces. Our pipeline would rather supplement the dHCP pipeline depending on the availability of T1 or T2 MRI data, since only one sequence may be available under the different clinical/research setting. In order to make our method more compatible with other group’s methods in terms of pial surface reconstruction and cortical thickness measurement, we aim to expand our pipeline to include T2w images in future studies.

Many neonates analyzed in our study were scanned twice, but each scan was treated as independent images when extracting their brain surfaces. While our approach is appropriate for cross-sectional analyses, past studies have proposed the segmentation/surface reconstruction methods that make use of information about the growth rate found in longitudinally collected MR images (Dai et al., 2013;Wang et al., 2013;Li et al., 2014;Nie et al., 2014;Li et al., 2015). This approach continues to be a research area of interest for future investigations aiming to improve cortical surface reconstruction.

Accurate reconstruction of neonatal brain WM and pial surfaces can be applicable to sampling cortical measurements of other modal MRI data which explain changes in different aspects of the brain. For example, diffusion tensor imaging is used to extract parameters related to WM integrity, such as fractional anisotropy and mean diffusivity. T1/T2 ratio myelin map (Glasser and Van Essen, 2011) is also shown to be a sensitive biomarker that predicts neonatal brain ages (Lewis et al., 2018) and cognitive outcomes (Moeskops et al., 2017). Using a boundary-based registration approach (Greve and Fischl, 2009) based on our reconstructed brain surfaces, we plan to compute these measurements within the cortex or on superficial WM, leading to more versatile applications that help clarify mechanisms involved in the early brain development.

Future studies also aim to elucidate the neurobiological and cellular basis for changes in cortical thickness trajectories. A possible explanation of such topological changes is through prematurity-related abnormalities in synaptic pruning, which refines developing neural circuits important for cognitive function and specialization (Knudsen, 2004;Raznahan et al., 2011b).

We have opened the NEOCIVET 2.0 to the public through CBRAIN (https://mcin-cnim.ca/technology/cbrain/), a web-based platform that distributes, processes, and exchanges 3D/4D brain imaging data. Our pipeline can also be used to investigate brain morphological changes in neonates born with clinical conditions.

## Acknowledgement

This study was supported by the National Institutes of Health grants (P41EB015922; U54EB020406; U19AG024904; U01NS086090; 003585-00001). HK was funded by Baxter Foundation Faculty Fellowship Award, and BrightFocus Research Grant award (A2019052S).

## Notes

### Competing Interest Statement

The authors have declared no competing interest.

