## supplementary materials for "Robust T1 MRI cortical surface pipeline for neonatal brain and systematic evaluation using multi-site MRI datasets"

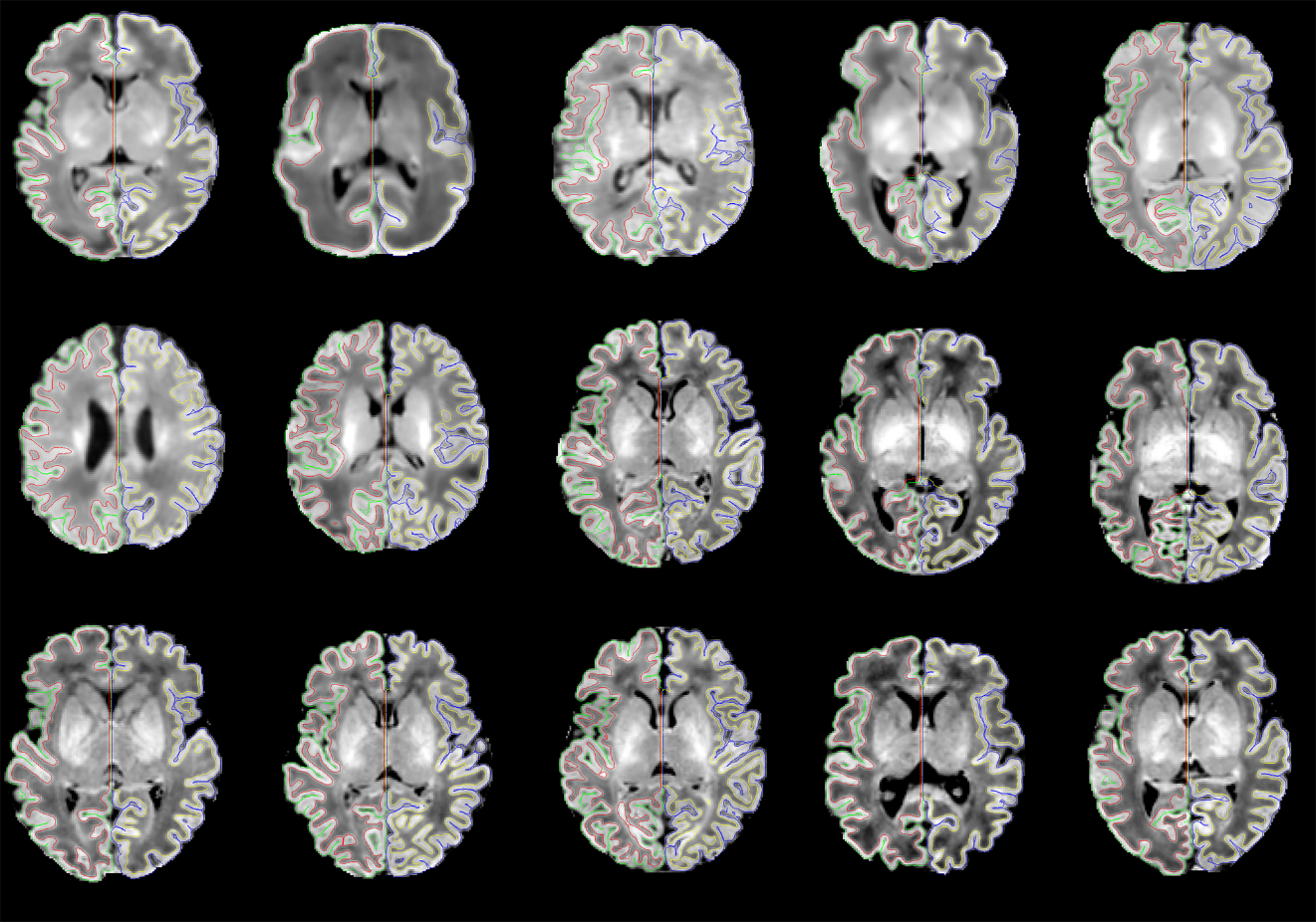

**Fig. S1**: illustrations of surface generated using our pipeline, with WM and pial surfaces overlaid. Images were randomly selected from all three datasets.

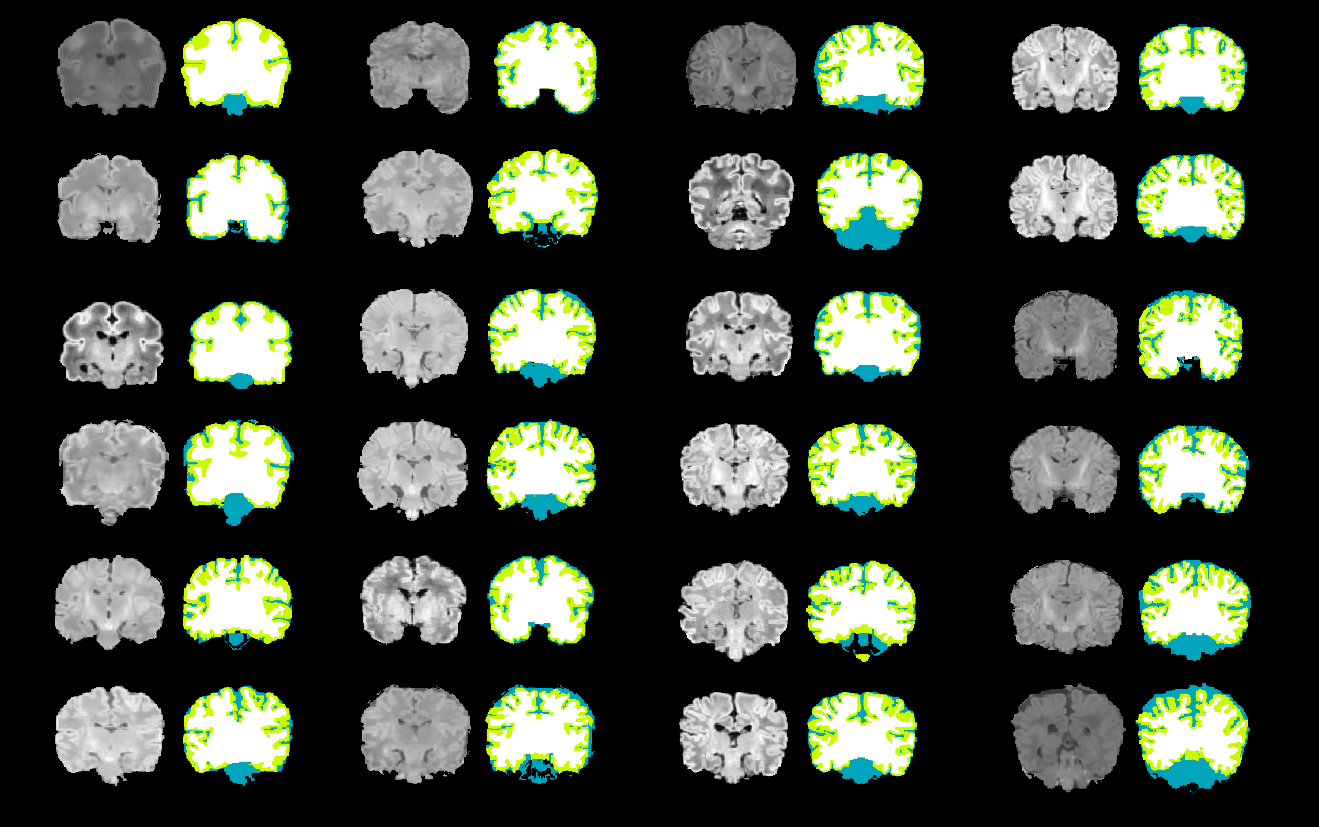

**Fig. S2**: All 24 intensity normalized and skull-stripped T1w image templates, together with the manual tissue segmentation used in our pipeline, the templates were placed with the order of ascending age.

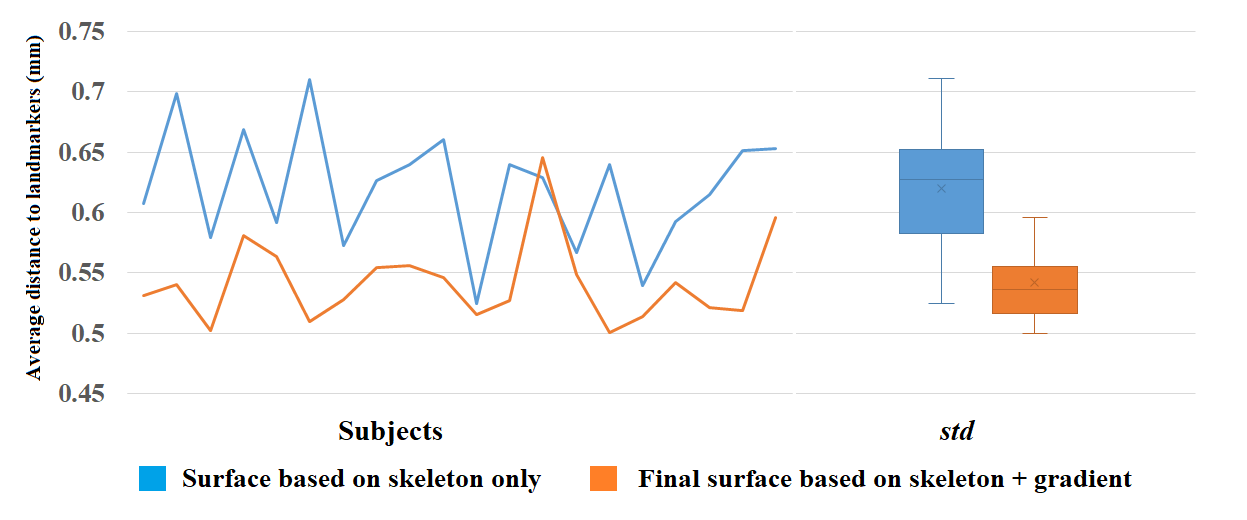

**Fig. S3.** Mean distance (mm) to the landmark from the CSF-skeleton (blue) and refined pial surface (orange) for the UCSF dataset. Subjects are sorted in terms of GA at scan (subject 1 is the youngest and subject 20 is the oldest).

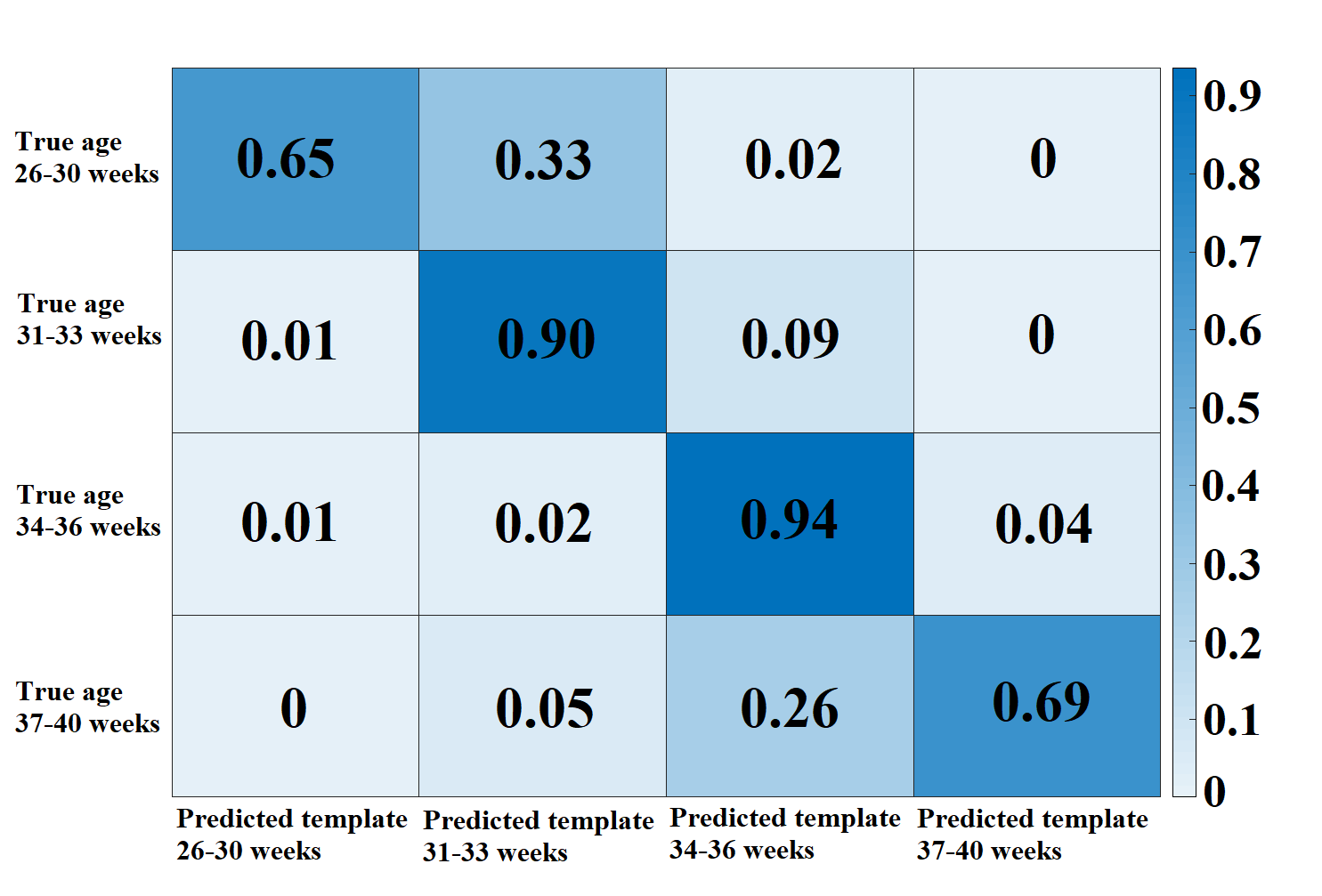

**Fig. S4.** Comparison of the choice of age-specific templates between using sulcal depth similarity and using postmenstrual age.

**Parameters selection**

Multiple steps associated with the brain extraction, tissue segmentation, skeletonization and surface reconstruction process required the selection of numerous parameters. All of the parameters introduced in parameter section are tested and operationalized in the images that are pre-processed and registered to the template space as explained in the manuscript. Thus, all of the images are normalized in brain size, orientation, spatial resolution and intensity distributions. On the other hand, the folding variations due to different maturations in relation to the gestational age and brain prematurity were addressed using the march-cube surface modeling. We thus believe that the influence of the choice of the same parameters on the accuracy of the surface extraction is minimal as shown in our qualitative and quantitative evaluations.

The parameters utilized in our pipeline are adopted empirically by visually checking the quality and convergence rate of the resulting surfaces. The adaptability of the parameters was cross-validated with three datasets from UCSF, dHCP and UNC. We delineated the important parameters and described their specific behaviors, such that others may apply our pipeline to their own dataset by using either verified default parameters from our three datasets or their own parameters customized from their own results.

a) *3D-Unet.* For both brain extraction and tissue segmentation, 3D-Unet was performed on patches instead of the whole T1w image volumes. Larger patches may decrease the number of training samples and increase the model parameters, which would cause the overfitting (insufficient memory would be another issue). Models trained on smaller patches may lose the ability to detect context information which in turn impair the segmentation. The patch size is recommeded to be set between 16 to 48. Overlap size for patches is also important in segmentation. The class label of each voxel is determined by majority-voting the labels predicted from different patches each voxel was involved in. Each voxel would be labeled more times when patch overlap was bigger, which may help increase the segmentation accuracy. However, insufficient memory would be a restriction for large overlap. Overlap size was recommended be set at least 50%, but no more than 80%, of the patch size.

b) *Skeletonization*. Gaussian curvature threshold, which determines the initial seeds for catchment basins, was set to -2. Users may select this value between -2 and 0. The mean curvature threshold plays an important role in erosion and ridge detector and was set to 0.1. Users should have this value no larger than 0.3. For the redundant branches pruning option, *curves* (including *border, curve, and curves* voxels*)* were set as the morphological pattern of branches that need to be truncated. Users may also set the pruning papameter to *interiors* which prunes the 6 outside neighbors of the *interior* voxelx, or the *interiors-curves* which prunes both *curves* voxels and the neighbors of *interior* voxels.

c) *Gradient refinement*. The gradient based surface refinement was performed in a *multi-resolution fashion* to efficiently fit the initial surface to the location displaying a regional maximum gradient. In each of the multi-resolution cycles, each set of the parameters including a number of blurring iterations, search distance (in mm), and search step (in mm) were empirically selected. We performed 4 cycles starting with the coarsest deformation and finishing with the finest deformation using the parameter sets: (90, 7.0, 0.7), (50, 5.0, 0.7), (20, 3.0, 0.3), and (20, 2.0, 0.2), respectively. We found that these sets were related to image characteristics including resolution, tissue contrast, and signal-to-noise, while not contingent upon the type of WM or pial surfaces. These sets were thus applied equally to both WM- and pial-surface refinements. Users can adjust their own parameters but the set of parameters has to be decreases in the 4 cycles.

**Comparison to dHCP pipeline**

A team in the developing human connectome (dHCP) project ([http://www.developingconnectome.org](http://www.developingconnectome.org/)) has most recently published a state-of-the-art toolkit to reconstruct neonatal inner and outer cortical surfaces on neonatal brain MRI (Makropoulos et al., 2018). There are several key technical differences between our pipeline and the dHCP pipline, as delineated in **Table S1**.

**Table S1**. Comparison between dHCP and our proposed pipelines

|  | dHCP | Proposed |
| --- | --- | --- |
| Neonatal scans tested | n= 411; PMA=29-45 weeks | n= 843; PMA=26-45 weekstested on three independent datasets (UCSF, UNC, dHCP) |
| Primary MRI modality | T2w | T1w |
| Brain extraction | BET | 3D-Unet |
| Tissue Segmentation | Draw-EM method | Non-local 3D-Unet |
| Model(s) of sulcal CSF course/location | Gradient edge of GM/CSF interface. | Skeletonization of CSF and GM union |
| Forces used in pial surface deformation | External force toward the sulcal GM/CSF edge  Non-self-intersection constraint  Smoothness constraint | External force toward GM/CSF Boundary defined as skeleton  Non-self-intersection constraint  Smoothness constraint |
| Pial surface deformation | Deform outward from the WM surface for a short distance followed by gradient-based refinement | Deform to skeleton-CSF union followed by gradient-based refinement |
| Multiresolution surface refinement | No | Yes |
| Surface template used to set point-wise correspondence | An average template constructed from term neonates | Age-specific templates |

Image processing in the dHCP pipeline is performed mainly on T2w images. Brain extraction is first performed on each subject's T2w image using FSL BET (Smith, 2002), and then the resulting brain mask is refined from Draw-EM segmentation. The reduced resolution and increased partial volume of neonatal data motivated the authors to use specially designed tissue segmentation (Makropoulos et al., 2012;Makropoulos et al., 2014;Makropoulos et al., 2016) and surface extraction protocols (Schuh et al., 2017). An edge-based surface refinement (gradient-based refinement similar to our approach) is applied to the WM surface generation. In regard to the process of pial surface deformation and refinement, the surface is first deformed outward for a short distance and then the refinement is performed to attract the surface to the GM/CSF boundary where the intensity gradient maximumly occurs, then the erroneous surface deformation is corrected by Laplacian smoothing, which smooths the distance between the deformed surface and the original surface. The constructed surfaces can be further registered to spatial-temporal cortical surfaces templates for between-subject comparisons (Bozek et al., 2016).

By contrast, our pipeline does not rely on T2w images and can be used with T1w images only, which are more ideal for general use. Neonatal brains exhibit dramatic morphological alterations throughout a wide range of gestational ages, which further compels us to perform brain extraction and tissue segmentation using deep learning-based algorithms (i.e. 3D-Unet for brain extraction and non-local 3D-Unet for tissue segmentation). Template samples are selected evenly from all gestational ages. To avoid severe partial volume effects, a skeletonization strategy is applied to construct a hypothetical GM/CSF interface. The WM surface is finalized by refining the intermediate surfaces generated from the WM segmentation. For the pial surface, the intermediate surface is first deformed to attach to the skeleton, then the gradient-based surface refinement is performed to finalize the pial surface deformation. Specifically, intensities at the GM/CSF boundary, where gradient maximumly occurs, are first recorded and corrected by Laplacian smoothing the intensities iteratively. Thereafter, the surface is deformed to fit the smoothed intensities. Finally, surface-based registration utilizing an age specific template was used to measure cortical gyrification, thereby capturing precise cortical morphology at specific ages which exhibit dramatic cortical folding changes throughout perinatal development.
